# Interplay between murine sex-biased gene expression and hepatic zonation revealed by single nucleus RNA sequencing

**DOI:** 10.1101/2022.01.18.476791

**Authors:** Christine N. Goldfarb, Kritika Karri, Maxim Pyatkov, David J. Waxman

## Abstract

The zonation of liver metabolic processes is well-characterized; however, little is known about the cell type-specificity and zonation of sexually dimorphic gene expression or its growth hormone (GH)-dependent transcriptional regulators. We address these issues using single nucleus RNA sequencing of 32,000 nuclei representing nine major liver cell types. Nuclei were extracted from livers from young adult male and female mice, from male mice infused with GH continuously to mimic the female plasma GH pattern, and from mice treated with TCPOBOP, a xenobiotic agonist ligand of the liver nuclear receptor CAR (*Nr1i3*). Analysis of these rich transcriptomic datasets revealed: **1)** expression of sex-biased genes and their key GH-dependent transcriptional regulators is primarily restricted to hepatocytes and is not a feature of liver non-parenchymal cells; **2)** many sex-biased transcripts show sex-dependent zonation within the liver lobule; **3)** gene expression is substantially feminized in both periportal and pericentral hepatocytes when male mice are infused with GH continuously; **4)** sequencing nuclei increases the sensitivity for detecting thousands of nuclear-enriched lncRNAs and enables determination of their liver cell type-specificity, sex bias and hepatocyte zonation profiles; **5)** the periportal to pericentral hepatocyte cell ratio is significantly higher in male than female liver; and **6)** TCPOBOP exposure disrupts sex-specific gene expression and hepatocyte zonation within the liver lobule. These findings highlight the complex interconnections between hepatic sexual dimorphism and zonation at the single cell level and reveal how endogenous hormones and foreign chemical exposure can alter these interactions across the liver lobule with large effects on both protein-coding genes and lncRNAs.

**Synopsis:** Single nucleus RNA-seq analysis elucidated the cell type-specificity and zonation of the sex-biased murine liver transcriptome, including thousands of long non-coding RNAs. Xenobiotic exposure induced widespread dysregulation, including both gain and loss of sex-biased gene expression and changes in zonation.

## Introduction

The liver is a sexually dimorphic organ characterized by the differential expression of hundreds of genes between males and females, including many cytochromes P450 and other metabolic enzymes [1]. The sex-differential expression of specific hepatic P450s and other drug-metabolizing enzymes [2] is a key factor in the sex dependence of liver metabolism of many drugs and steroids in both rodents [3–5] and in humans, where sex-biased drug metabolism and pharmacokinetics is common [6, 7]. Sex-differences also characterize the incidence and severity of major liver diseases, including non-alcoholic fatty liver disease, steatohepatitis and hepatocellular carcinoma, which are more prevalent and more severe in males [8–10]. A hormonal basis for these sex differences is indicated by the lower rates of disease in premenopausal females as compared to males and post-menopausal females [11].

Liver sex differences are largely determined by the distinct temporal patterns of pituitary GH secretion between the sexes. In males, GH is secreted in discrete pulses of hormone release, followed by periods when GH is essentially undetectable in circulation; whereas in females, pituitary GH secretion occurs more frequently, resulting in a near-continuous plasma GH profile [12–14]. In males, each plasma GH pulse initiates a wave of liver GH receptor signaling via the JAK-STAT pathway [15, 16], most notably tyrosine phosphorylation to induce dimerization and nuclear translocation of the GH-responsive transcription factor Stat5b, followed by transcriptional activation of its downstream gene targets [1]. In female liver, persistent (near continuous) GH signaling partially down regulates Stat5b activity [17] via feedback inhibition of the overall signaling pathway by factors such as Socs2, a primary target gene of Stat5b [18]. GH-activated Stat5b is an essential transcriptional regulator of liver sex-biased genes, with Stat5b positively regulating up to 90% of male-biased genes while repressing 60% of female-biased genes in male mouse liver [19, 20]. Other transcription factors essential for establishing and/or maintaining liver sex differences include Hnf4a [21, 22], Hnf6, Cux2 [23, 24] and Bcl6 [25–29]. The crucial role of plasma GH profiles in regulating liver sex differences is further supported by the substantial feminization of the liver seen when male mice are given a continuous infusion of GH (cGH) over several days [22, 30, 31] and by the partial restoration of male gene expression patterns when hypophysectomized mice are given exogenous pulses of GH [32].

GH and Stat5b regulate many liver-expressed long non-coding RNAs (lncRNAs) in a sex-dependent manner, a subset of which are altered in expression following hypophysectomy or continuous GH infusion in male mice [33–35]. Many such lncRNAs have nuclear regulatory functions [36] and are tightly bound to liver chromatin [35]. LncRNAs show high tissue and cell type specificity [37, 38], consistent with their diverse roles in complex biological systems, including the liver, where they impact disease development [39–41] and present therapeutic opportunities [42]. LncRNAs can impact specific liver cell-type populations, for example, the lncRNAs Hottip and Meg3, which respectively activate and inhibit hepatic stellate cell activation [43, 44]. Expression patterns have been reported based on bulk RNA-seq analysis for more than 15,000 liver-expressed lncRNAs under a variety of conditions, including differences in sex [33–35] and foreign chemical exposure [35, 45–47], however, the liver cell type specificity of the vast majority of liver-expressed lncRNAs is unknown.

Hepatic stresses induced by high fat dietary regimens [48, 49] and chemical exposures [50–52] are often sex-dependent, are modulated by GH signaling [53, 54], and are associated with the development of liver diseases [11, 55]. Furthermore, many hepatic stresses induce pathologies within the liver lobule in a zone-dependent manner [56, 57], making it important to determine whether sexually dimorphic genes and functions can be ascribed to specific liver cell types or zonal regions of the liver lobule. The liver lobule is metabolically zonated, with pathways such as gluconeogenesis and ureagenesis enriched in hepatocytes of the periportal region, supported by proximity to fresh blood supply of oxygen and nutrients, while pericentral hepatocytes are enriched for glycolysis and xenobiotic metabolism [58, 59]. Single cell (sc)RNA-seq has been used to characterize zone-dependent gene expression across the liver lobule in hepatocytes [60], liver endothelial cells [61] and in hepatic stellate cells [62], linking physiological and pathological functions to zonated liver cell sub-populations. However, the impact of this zonation on sex-biased gene expression, including lncRNA expression, has not been defined. Moreover, it is unclear whether hepatic zonation itself may differ between the sexes.

Here, we characterize the transcriptomes of 32,000 individual nuclei extracted from male and female mouse livers, including livers from males infused with GH continuously for one week, which substantially feminizes hepatic gene expression [31], and livers from mice exposed to the CAR agonist ligand TCPOBOP, which dysregulates hundreds of genes, including many lncRNA genes [45, 46]. Our findings establish that sex-biased gene expression in the liver is largely limited to hepatocyte populations, and that many sex-biased transcripts are zonated within the liver lobule, with a subset showing sex-dependent zonation. Further, we show that single nucleus (sn)RNA sequencing is highly sensitive for detection of nuclear transcripts and can be used to detect thousands of liver-expressed lncRNAs, including sex-biased lncRNAs, and then establish their cell-type specificities and zonation bias. Overall, our findings highlight the interconnectedness of liver sexual dimorphism and hepatic zonation and illustrate how these processes interact to regulate the complex sexually dimorphic patterns of gene expression in the liver.

## Results

### Overview of snRNA-seq workflow

Frozen liver tissue pooled from four livers per group was used to generate single nuclei for each of 6 different control and treatment groups, representing five different biological conditions: male and female mouse liver, livers from male mice treated with continuous GH (cGH) for 7 days and a corresponding untreated male control, and TCPOBOP-treated male and female mouse liver. Livers were homogenized in a salt/detergent-based lysis buffer, nuclei were pelleted at low speed, washed in the presence of RNase inhibitors then passed through a strainer to remove aggregated nuclei and cell debris (**Fig. S1A**). Single nuclei were captured in gel bead-in-emulsions and barcoded using 10x Genomics Chromium instrumentation followed by Illumina sequence library preparation and high throughput RNA sequencing.

Sequencing datasets from all 6 liver snRNA-seq libraries were normalized and integrated after removal of doublets and high mitochondrial content nuclei after correction for inter-sample and inter-batch variation, resulting in a single harmonized data set comprised of ∼32,000 nuclear transcriptomes. These nuclei were clustered as a single group and projected in two dimensions on a UMAP (**Fig. 1A**), where nuclei with similar transcriptomes are found nearby each other [63].

**Fig. 1.**
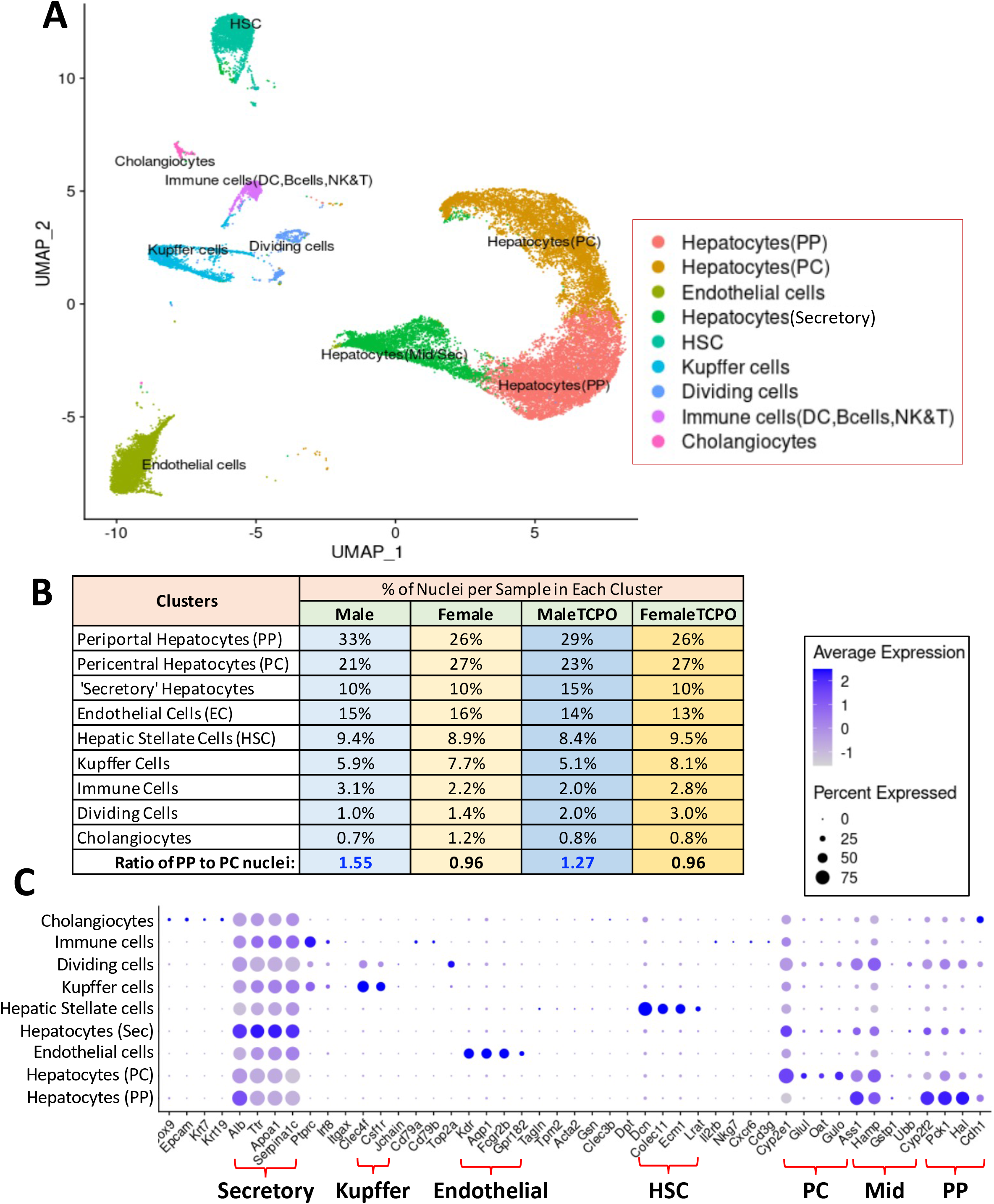
UMAP Clusters of liver nuclei identified by cell-type specific markers. 31,896 nuclei from all six liver samples (**Table S1B**) were clustered into 9 cell-type clusters after removal of cell multiplets and nuclei with high mitochondrial content. (**A**) UMAP representation of the 9 clusters. (**B**) Distribution of nuclei across the 9 clusters as a percentage of the total number of nuclei in each cluster. Also see **Table S1B**. (**C**) Dot plot expression for marker genes listed on x-axis. Dot color, average expression level of the gene in the cluster; dot size, percentage of nuclei expressing the gene. Cluster identity was based on established cell type-specific markers from literature. PP, periportal; PC, pericentral; Mid, midlobular.

### Liver cell cluster characterization

We identified three main hepatocyte clusters, comprising ∼65% of all 32,000 nuclei, a small cluster with hepatocyte and dividing cell characteristics, and five major NPC clusters (**Fig. 1, Table S1B**). The two largest hepatocyte clusters were identified as periportal and pericentral hepatocytes based on their marker genes (**Fig. 1C**), and the largest NPC clusters were endothelial cells (marker gene, Kdr), hepatic stellate cells (Dcn), and Kupffer cells (Clec4f), each comprising 5-20% of all liver nuclei. Smaller NPC clusters were identified as cholangiocytes (epithelial cells of the bile duct; marker gene Epcam) and immune cells (Ptprc), each comprising 0.7-3.9% of all liver nuclei. Kupffer cells were more abundant in female than male liver, as was reported in the rat [64, 65], both with and without TCPOBOP exposure (**Table S1B**).

The pericentral hepatocyte cluster was subdivided into two sub-clusters at higher resolution (**Table S1C**). The cluster tip region was enriched for highly pericentral-specific transcripts, most notably Glul, Slc1a2, and the lncRNA Meg3/lnc10922, an inhibitor of liver fibrosis [66], while the main cluster body, which bordered the periportal region, displayed mid-lobular characteristics. Another large cluster, positioned adjacent to the periportal cluster in the UMAP, expressed a mixture of pericentral, mid-lobular and periportal marker genes at moderate levels, with no clear enrichment for any zonation markers compared to periportal or pericentral hepatocytes. Hepatocyte-specific secretory protein RNAs, such as Apoe, Hp and Saa1, showed up to 4-5-fold enrichment in this cluster as compared to the other hepatocyte clusters (**Fig. 1C, Table S1D**). This cluster, which is not seen in mouse liver scRNA-seq datasets [60, 67], is likely comprised of hepatocyte nuclei associated with membrane-bound polyribosomes, which contain high levels of mature (spliced) RNAs for abundant liver secretory proteins.

We identified a small cluster characterized by the dividing cell marker Top2a, a gene involved in DNA replication [68]. These dividing cell nuclei were 40-50% more frequent in TCPOBOP-treated livers (**Fig. 1B**), where hepatocyte cell division is increased [69, 70]. This cluster was identified as dividing hepatocytes based on its enrichment for hepatocyte-specific genes, primarily those from pericentral and mid-lobular hepatocytes, when compared to NPCs (**Fig. 1C** and **Fig. S1D; Table S1E**). Confirming this, DAVID functional annotation analysis identified oxidoreductase/cytochrome P450 metabolism, peroxisomes and nuclear receptors as major enriched functions of the dividing hepatocytes (**Table S1E**).

We observed a higher periportal to pericentral cell ratio in male liver (1.55:1) than female liver (0.96:1) (**Fig. 1B**), which may reflect differences in physiological requirements between the sexes. A higher frequency of periportal hepatocytes in males was also seen in an independent single cell dataset from the Tabula Muris study [67] (**Table S1B**), and when comparing TCPOBOP-treated male vs female liver (**Fig. 1B**). The same trend was seen when comparing nuclei from a second batch of male liver to nuclei from male mice given GH as a 7-day infusion (cGH), which feminizes the overall pattern of hepatocyte gene expression (see below), and correspondingly, decreased the fraction of periportal hepatocytes substantially (**Table S1B**).

### Sex-biased gene expression is largely restricted to hepatocytes

Genes showing sex-biased expression were identified by male vs female differential expression analysis applied to each cluster. A total of 1787 genes showed sex-biased expression in one or more of the 9 clusters examined at FDR < 0.05. Remarkably, 1759 of the 1787 genes (98%) showed sex-biased expression in hepatocytes (**Table S2A, Table S2B;** 784 protein-coding genes, 975 lncRNAs). Moreover, for 1669 of the 1787 genes (93%), sex-biased expression was restricted to hepatocytes. In contrast, only 28 genes (1.6%) showed sex-biased expression restricted to one of the five NPC clusters: endothelial cells (13 genes), hepatic stellate cells (10 genes), and Kupffer cells (5 genes) (**Table S2B**). Five genes, including two Y-chromosome genes, showed male-biased expression in all 9 clusters, and correspondingly for 6 female-biased genes. The latter genes includes the lncRNA Xist (lnc15394), which showed strong female-biased expression (> 370-fold) in all clusters (**Fig. 2A**), as was expected due to its widespread role in X-chromosome dosage compensation in female eutherian mammals [71]. Genes with strong sex-biased expression restricted to hepatocyte clusters include the male-biased Cyp4a12, the female-biased Slc22a26 (**Fig. 2B-2C**), and many sex-specific lncRNAs and other genes (**Fig. S1E**).

**Fig. 2.**
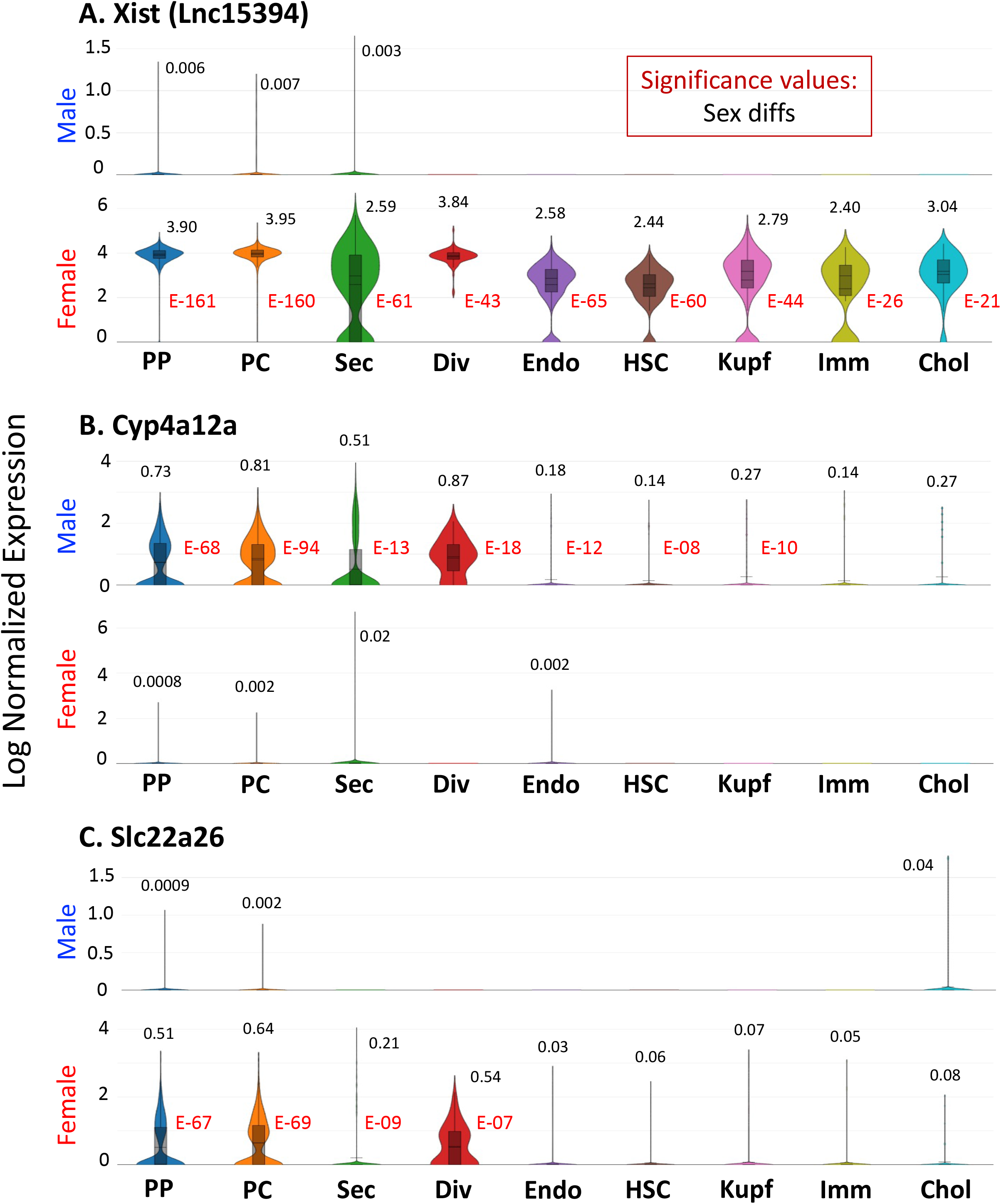
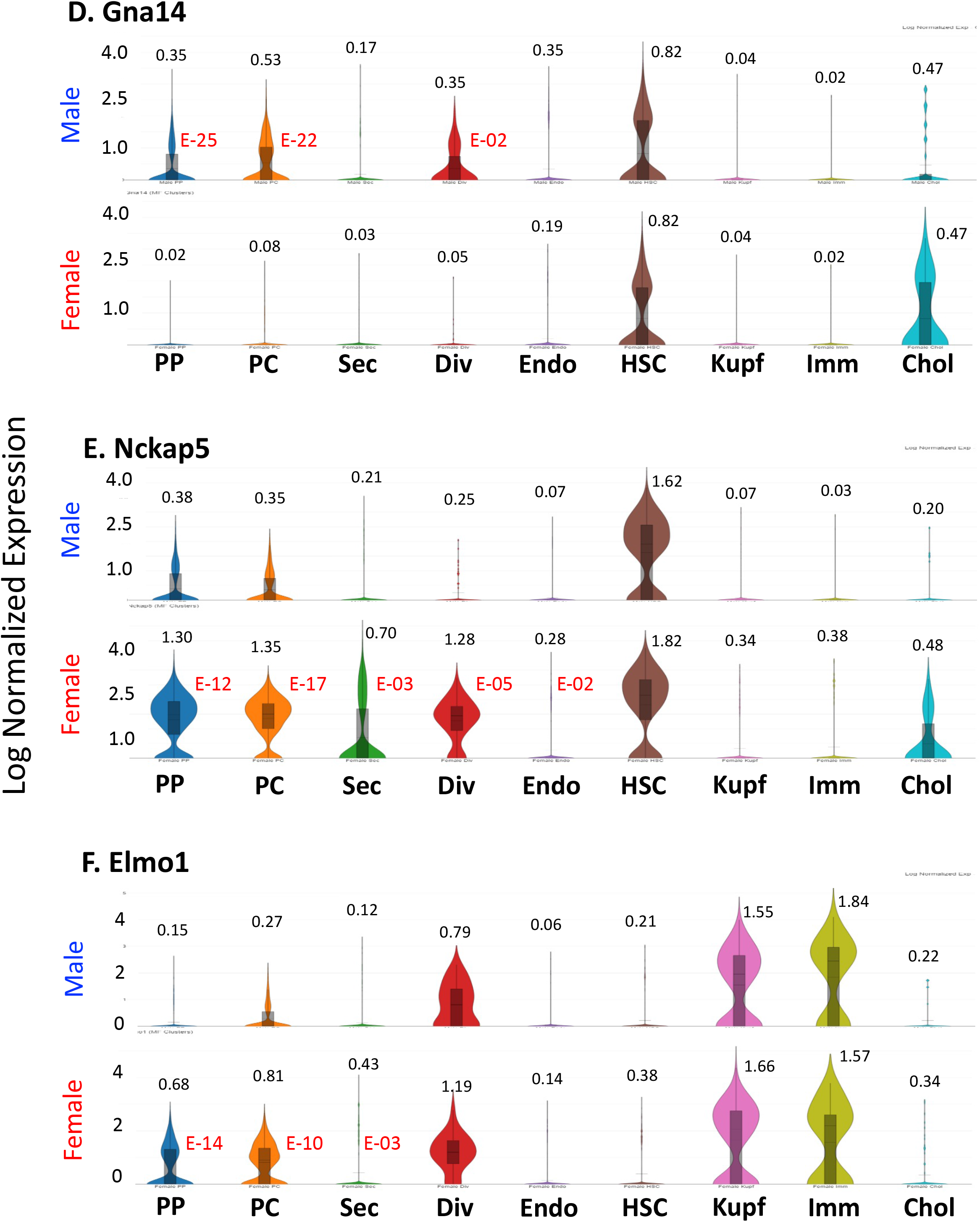
Expression of select sex-biased genes in 9 liver nuclei clusters from male and female control mouse liver. (**A**) snRNA-seq data for lnc15934 (Xist), which shows strong sex-biased expression in nearly all cell-type clusters, (**B**) for Cyp4a12a, and (**C**) for Slc22a26, which respectively show male-biased and female-biased expression in hepatocytes only. (**D-F**) Genes that show sex-biased expression in hepatocytes, but show non- sex-biased expression in NPCs. Violin plots indicate the distribution of individual nuclei-based cell barcode UMI counts for each cluster. Average UMI count expression values for each cluster, log normalized across all nuclei in all clusters for that sample. Red numbers to the right of each violin plot represent fold change and significance (FDR) values for differential expression when comparing the pericentral and periportal hepatocyte clusters, taken as a single group, to the five NPCs clusters, also taken as a single group (**Table S2E**). NS, not significant. Cell clusters are as marked at the bottom: PP, periportal; PC, pericentral; Sec, secretory hepatocytes; Div, dividing cells; Endo, endothelial cells; HSC, hepatic stellate cells; Kupf, Kupffer cells; Imm, immune cells; Chol, cholangiocytes.

While many of the sex-biased genes identified here were previously identified by bulk liver nuclear RNA-seq analysis, many others were novel (**Table S2B**). Discovery of these novel sex-biased genes reflects the increased sensitivity provided by snRNA-seq. In some cases, this increased sensitivity results from the exclusion from the hepatocyte clusters of sequence reads derived from NPCs that express the gene at a comparatively high level but without sex bias, as shown for Gna14 and Nckap5 in HSCs (**Fig. 2D**, **Fig. 2E**) and for Elmo1 in Kupffer cells and Immune cells, as well as Dividing hepatocytes (**Fig. 2F**). In addition, by sequencing liver nuclei, rather than liver cells, we greatly increased the sensitivity for detection of the many chromatin-bound lncRNAs that are sex-biased [35]. Consequently, we identified many fewer sex-biased lncRNAs in our analysis of the Tabula Muris mouse hepatocyte single cell RNA-seq dataset, a widely used reference [67].

### Hepatocyte specificity of GH signaling regulators and transcription factors

Liver sex differences are regulated by the distinct, sex-specific pituitary GH secretion patterns: pulsatile secretion in males, and persistent/near-continuous secretion in females. We examined three key genes of GH signaling that are essential for liver sex differences [19]: GH receptor (Ghr), Jak2, a tyrosine kinase that is activated by Ghr, and Stat5b, a transcription factor activated by Jak2-catalyzed tyrosine phosphorylation. We also examined Socs2, a feedback inhibitor of GH receptor signaling. Expression of Ghr, but not the other factors, was significantly higher (FDR = 2.1E-7) in hepatocytes as compared to the 5 NPC clusters (**Fig. S2A-S2B**). Jak2 expression was low in all clusters, and Stat5b and Socs2 were more highly expressed in hepatocytes than NPCs, albeit not significantly.

We also examined 15 liver transcription factors, most of which were readily detected by snRNA-seq in one or more clusters (**Fig. S2C-S2F**). Highest expression was typically found in periportal and pericentral hepatocytes (and in dividing hepatocytes), where many of the factors were significantly enriched compared to the 5 NPC clusters, in either male or female liver. Of note, seven of the transcription factors exhibited significant sex-bias in their expression (**Fig. 3**), in agreement with prior findings in bulk liver RNA-seq and other studies [18, 23–25, 31, 72]. The strong hepatocyte-bias of these sex-dependent, liver-enriched transcription factors is likely key for the hepatocyte-specificity of sex-biased gene expression. Many other novel sex-biased transcription factors and transcriptional regulators were identified in the full dataset, with 102 of the 1787 sex-biased genes containing the Gene Ontology term ‘transcription’ (**Table S2**, column AB).

**Fig. 3.**
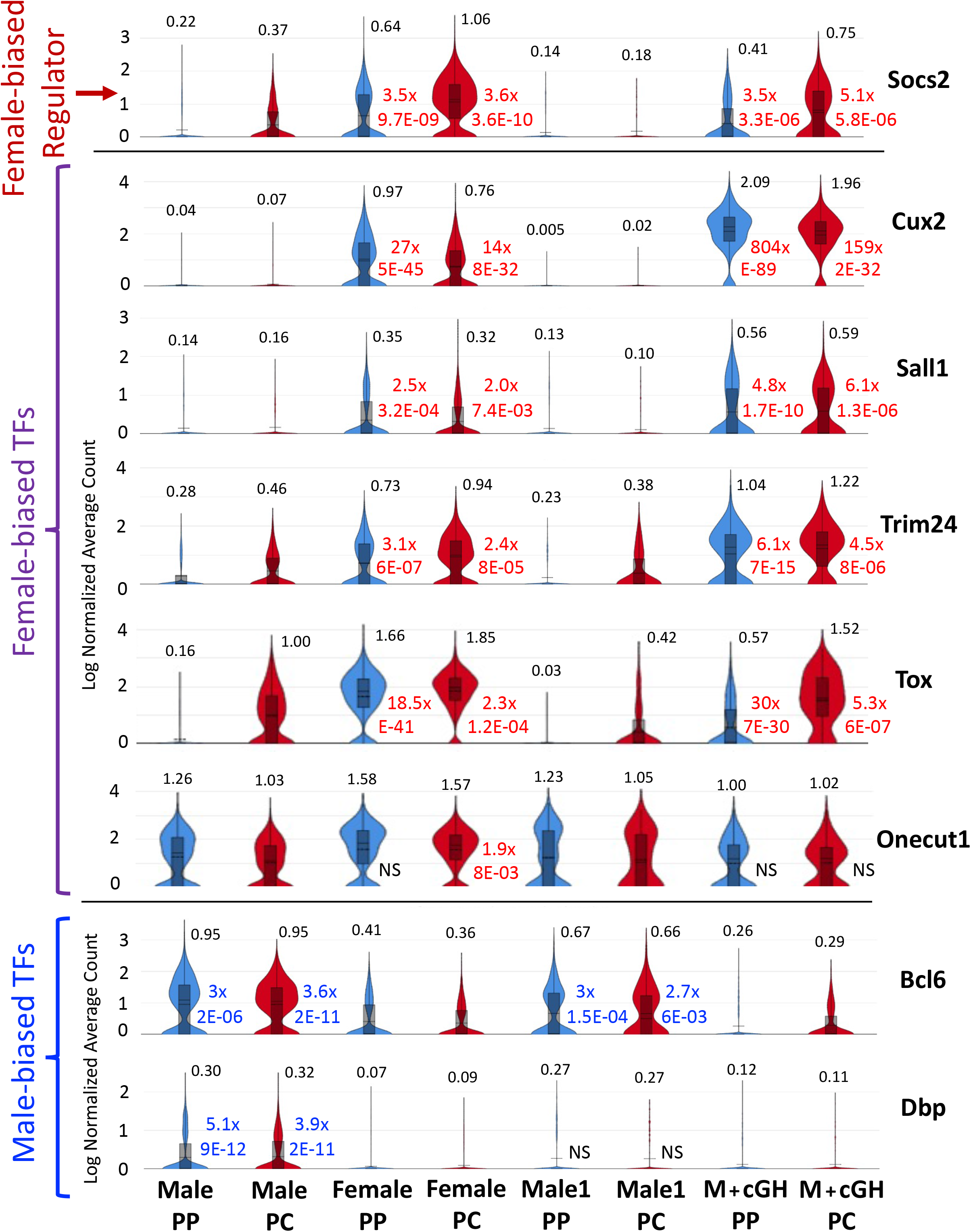
Expression of sex-biased transcription factors (TFs) and Socs2 in male and female liver and in cGH- treated male liver. Violin plots (as described in Fig. 2) showing expression data for the female-biased GH signaling inhibitor Socs2, for 5 female-biased transcription, and for 2 male-biased transcription factors in both periportal (PP) and pericentral (PC) hepatocyte clusters from male and female liver (**Table S2B**), and from male control (Male-1) and cGH-infused male liver (**Table S4A**). Data presentation as in Fig. 2, except that values to the right of each violin plot (red or blue) indicate fold change values (expression ratios) and significance for sex bias (first four violins in each row) or for the response to cGH infusion (last four violins) for comparisons between the two specific cell clusters being compared, i.e., Female PP vs Male PP, Female PC vs Male PC, Male+cGH PP vs Male-1 PP, and Male+cGH PC vs Male-1 PC, where Male1 data are from the Male-1 controls for the cGH-treated male group.

### Expression of sex-biased genes in periportal versus pericentral hepatocytes

Many genes showed sex-biased expression in both periportal and pericentral hepatocytes. Examples include lnc5999 (>39-fold female bias) and Nox4 (>15-fold male bias), whose strong, sex-biased expression was seen in both hepatocyte populations (**Fig. 4A**, **Fig. 4B**). However, other sex-biased genes showed strong, sex-biased expression in only one of the two major hepatocyte clusters. For example, lnc18328 was highly expressed with a strong female bias (>1000-fold) in pericentral hepatocytes but not in periportal hepatocytes, where its expression was reduced >97% (**Fig. 4C**). Chrm3, which plays a role in liver injury response [73] and showed strong male-bias (10-fold) in periportal hepatocytes, gave 85% lower expression in pericentral hepatocytes (**Fig. 4D**). Sex-biased genes such as these are likely under the control of both sex-dependent regulators and zonation regulatory factors. Transcription factors that show zonated, sex-biased expression and that may contribute to the zone-dependence of sex-biased gene expression include Tox, a female-biased transcription factor with a much higher female bias in periportal than pericentral hepatocytes (18.5-fold vs 2.3-fold; **Fig. 3, Table S2B**).

**Fig. 4.**
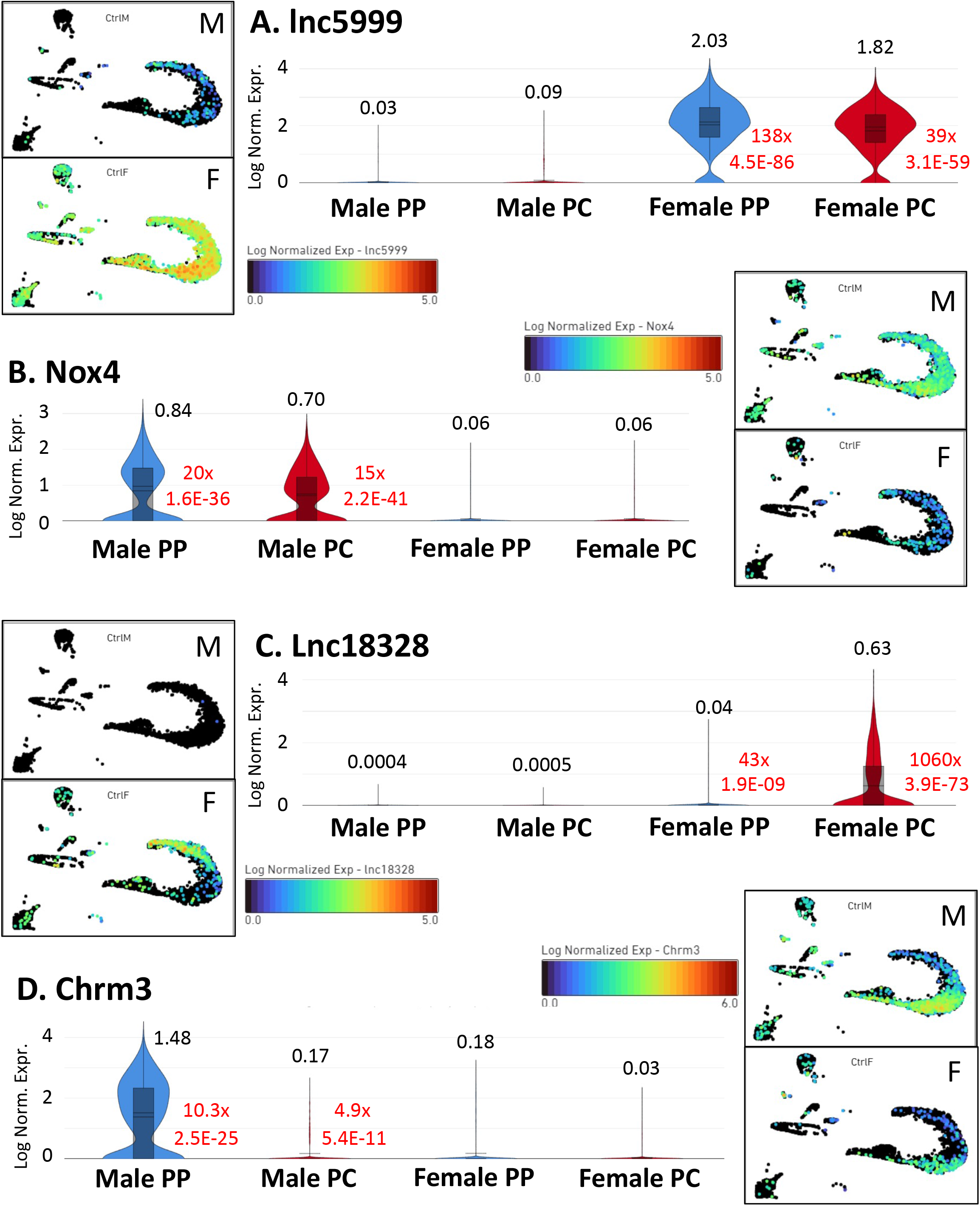
Expression of sex-biased genes in periportal and pericentral hepatocytes. Expression data for genes showing strong sex-biased expression in both periportal (PP) and pericentral (PC) hepatocytes (**A, B**), and for genes whose sex-biased expression is much stronger in pericentral hepatocytes (**C**) or in periportal hepatocytes (**D**). Sex-biased expression data is from **Table S2B**. Images alongside each panel are UMPAs (as in Fig. 1A) indicating log normalized expression in each nucleus in male (M, upper image) and female liver (F, lower image) per the color bar legend for each gene. Data presentation as in Fig. 2, except that red values to the right of each violin plot indicate fold change values (expression ratios) and significance for Male-Female differential expression in periportal or pericentral hepatocytes.

Next, we directly evaluated hepatocyte zonation-bias in both male liver and female liver. We identified 719 genes, 376 protein-coding genes and 343 lncRNAs, that show zonated expression in at least one sex (**Table S3A**). We found a similar number of zonated protein-coding genes but 8-12-fold fewer zonated lncRNAs when using the scRNA-seq Tabula Muris data set [67] (**Table S3G**), consistent with the lower sensitivity of scRNA-seq for detection of lncRNAs. Cyp7a1 is an example of a gene with strong pericentral-biased expression in both sexes (pericentral/periportal expression ratio = 6-15) (**Fig. 5A**), while Sds shows strong periportal bias (periportal/pericentral = 7) in both sexes (**Fig. 5B**). The full set of periportal-biased genes was enriched for histidine metabolism and carbon metabolism, while pericentral-biased genes were enriched for cytochrome P450 and lipid metabolism and ion transport (**Table S3C**, **Table S3D**), consistent with earlier work [74, 75] and validating these two major hepatocyte subpopulations.

**Fig. 5.**
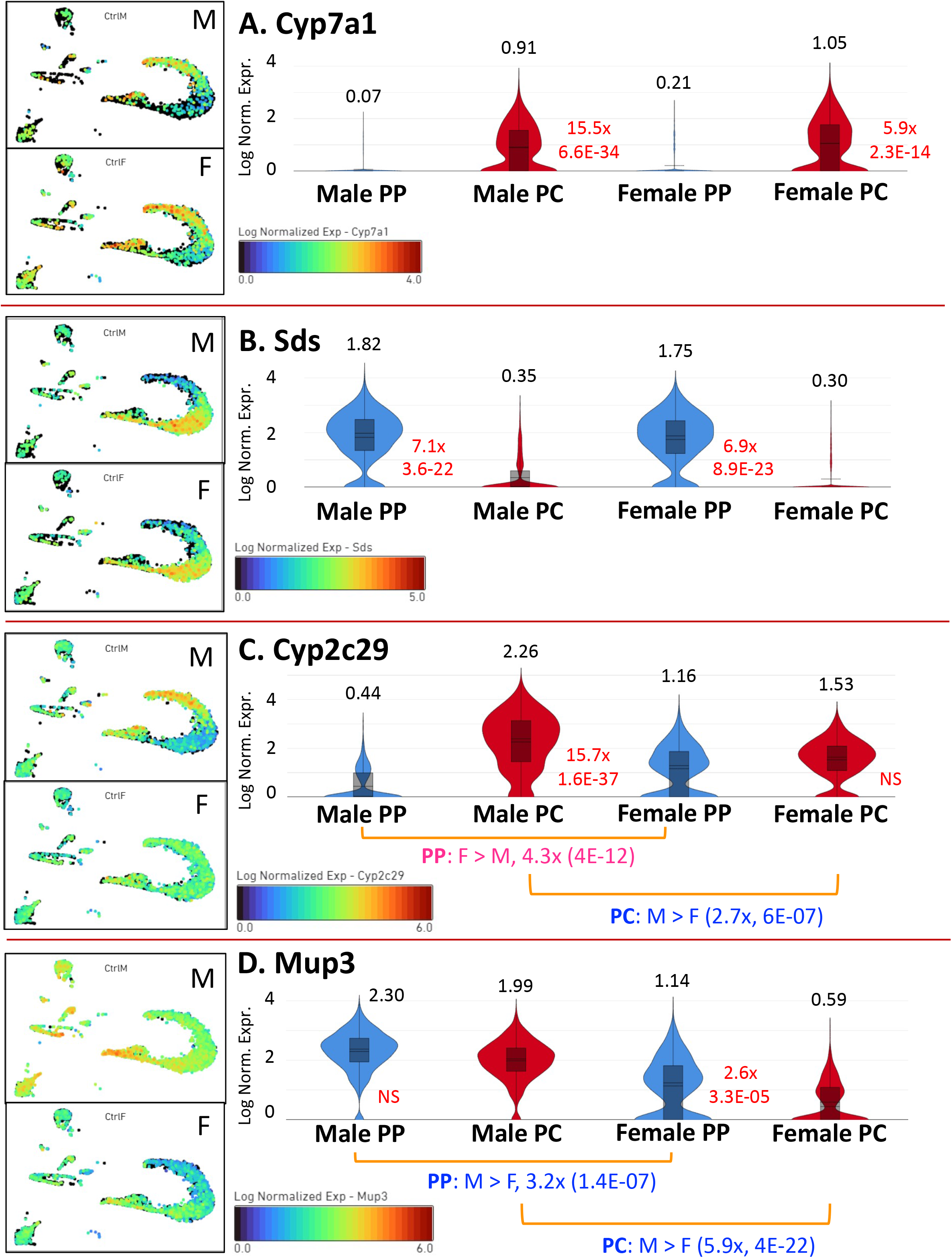
Zonation-biased expression in male and female liver. Expression data for genes showing significant zone-dependent expression in both male and female liver (**A, B**), only in male liver (**C**), or only in female liver (**D**), based on expression data from **Table S3A**. UMAP images display the log normalized expression in each nucleus in male (upper image) and female liver (lower image), per the color bar legend for each gene. Data presentation as in Fig. 2, except that red values indicate fold change values (expression ratios) and significance for periportal versus pericentral expression in each sex. Sex differences are marked below each panel in (**C**) and (**D**).

Strikingly, 333 (46%) of the 719 zonation-biased genes are sex-biased genes (**Fig. S3**), highlighting the interconnection of these two mechanisms of gene regulation. Many of these genes showed zonation-biased expression in only one sex (**Fig. S4, Table S3B**), which in some cases could simply be due to expression in one sex being too low to meet our thresholds for significant zonation. However, in many cases we observed a clear sex-dependence to zonation (**Table S3A, column J**). The most striking example is Cyp2c29, whose pericentral bias in males appears to drive its sex-differential expression in both hepatocyte populations (**Fig. 5C**). Thus, Cyp2c29 is 15.7-fold pericentral-biased in male liver, but in female liver it shows no significant zonation. This zonation drives Cyp2c29 to be male-biased pericentrally but female-biased periportally by a mechanism that most likely involves a combination of zonation and sex-biased regulators. Mup3, which shows significant male-biased expression in both hepatocyte populations, is another interesting example where zonation bias is limited to one sex. Specifically, Mup3 shows a 2.6-fold periportal bias in expression in female liver but no zonation bias in male liver (**Fig. 5D**). This pattern is consistent with the proposal that in male liver, sex-biased transcription factors activate Mup3 to its maximum level in both hepatocyte populations, overriding the influence of any zonation factors, while in female liver zonation factors dictate the observed periportal bias in expression. Together, these findings highlight a complex and diverse set of interconnected regulatory mechanisms controlling sex-biased zonation and zonation-biased sexual dimorphism in hepatocytes.

### Feminization of hepatocyte gene expression by GH infusion

Continuous infusion of GH in male mice overrides endogenous male plasma GH pulses and significantly, albeit incompletely feminizes sex-biased liver gene expression within one week [31]. However, it is not known whether both periportal and pericentral respond equally well to cGH infusion. A1bg is a classic example of a female-biased gene that we found is strongly up-regulated in male liver by cGH infusion (>400-fold increase) in both hepatocyte populations (**Fig. 6A**), while Cyp2d9 is a male-biased gene that is strongly down-regulated by cGH, more completely in periportal than pericentral hepatocytes (42-fold vs. 9.4-fold decrease; **Fig. 6B**). Female hepatocyte gene expression patterns were also recapitulated in cGH-infused male liver for Socs2 and for 5 of the 7 sex-biased liver transcription factors described above (**Fig. 3**, right four columns). By feminizing the expression of these sex-specific regulators, cGH feminizes many downstream sex-biased genes (**Table S4A**). Thus, by overriding the effects of the endogenous plasma GH pulses in male mice, cGH reprograms the expression of 547 sex-biased genes, including 290 sex-biased lncRNAs, in either periportal or pericentral hepatocytes from male liver (**Tables S4A-S4B**). Overall, 246 of 285 cGH-responsive female-biased genes were up regulated in male liver, while 253 of 261 cGH-responsive male-biased genes were down-regulated (Fisher extract test, p < 0.0001) (**Table S4B**).

**Fig. 6.**
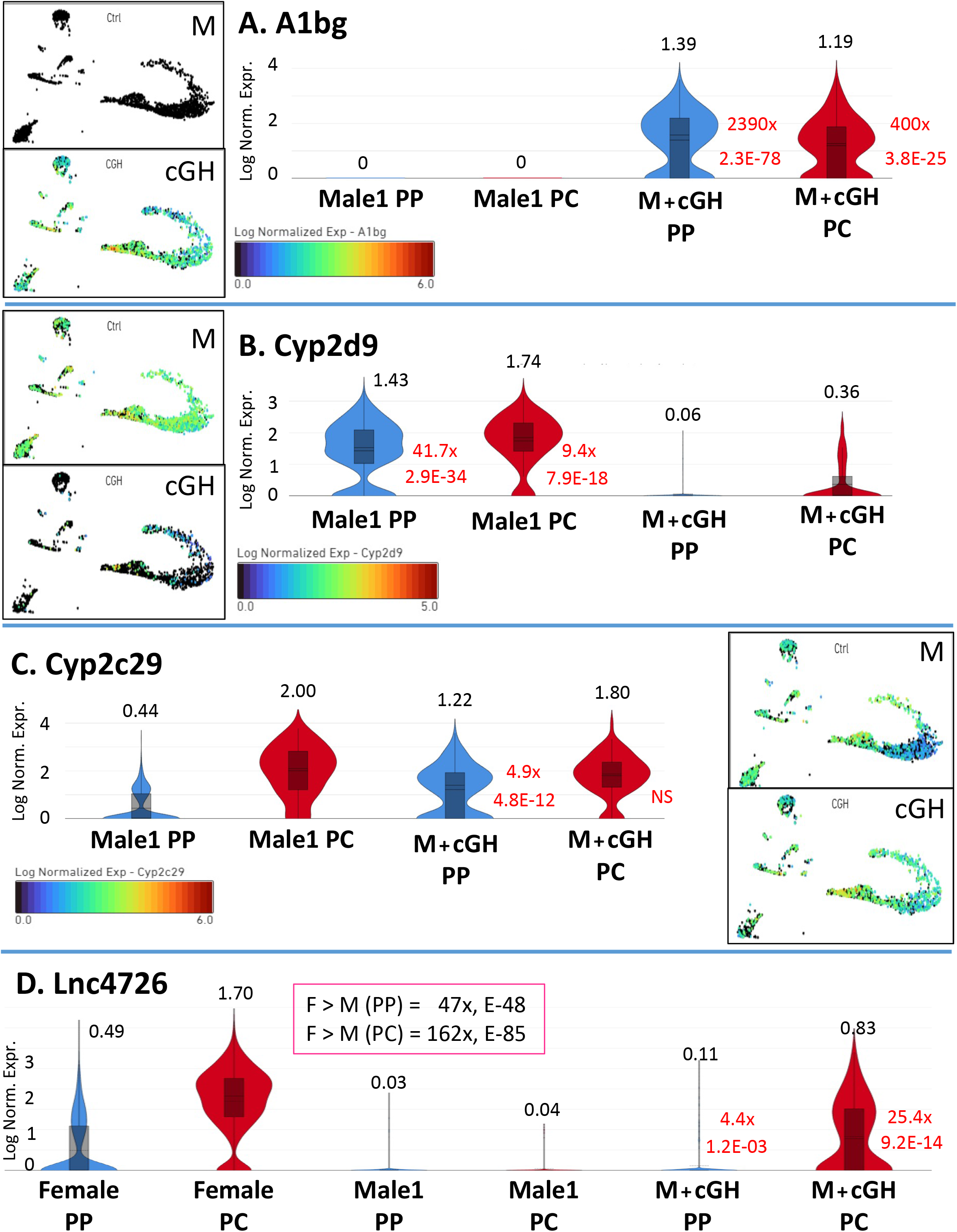
Impact of cGH infusion on expression of sex-biased genes in male liver. Expression data (**Table S4**A) for sex-biased genes whose induction (**A,** A1bg) or repression (**B,** Cyp2d9) following cGH infusion for 7 days is significant in both periportal (PP) and pericentral (PC) hepatocytes. Correspondingly, A1bg shows strong female-biased expression and Cyp2d9 shows strong male-biased expression in both hepatocyte populations (**Table S2B**). Also shown are sex-biased genes whose induction by cGH in male liver primarily occurs in periportal hepatocytes (**C,** Cyp2c29), or in pericentral hepatocytes (**D,** Lnc4726). Cyp2c29 shows significant female-biased expression in periportal hepatocytes but male-biased expression in pericentral hepatocytes. Lnc4726 shows significant female-biased expression in both hepatocyte clusters, but with significant pericentral zonation in female liver (4.4x, E-10; **Table S3A**), consistent with the high pericentral induction by cGH seen here. UMAP images display log normalized expression in each nucleus in control male (top) and in cGH male-infused liver (bottom), as per the color bar legend for each gene. Data presentation as in Fig. 2, except that red values indicate fold change values (expression ratios) and significance for the response to cGH infusion in that cell cluster. Male1, data are from the Male-1 controls for the cGH-treated male group.

### Hepatocyte zone-specific responses to cGH infusion

A substantial subset of the cGH-responsive sex-biased genes, comprised of 283 genes, responded to cGH infusion in only one of the two major hepatocyte clusters (**Fig. S5, Table S4B**). One striking example is Cyp2c29, which showed strong pericentral bias in expression in untreated male liver (**Fig. 5C**) and was strongly induced by cGH in periportal hepatocytes but showed no significant response in pericentral hepatocytes (**Fig. 6C**). Thus, cGH infusion not only abolishes the strong female bias in periportal Cyp2c29 expression seen in untreated mice, but also the zonal difference in expression seen in male but not female mouse liver. Thus, in male periportal hepatocytes, cGH overrides the repressive actions of plasma GH pulses to induce Cyp2c29 expression (4.9-fold, FDR = 4.82 E-12), whereas in pericentral hepatocytes Cyp2c29 shows no response to cGH (**Fig. 6C**). Lnc4726 exemplifies a highly female-specific gene with a strong pericentral bias. cGH infusion induces this lncRNA > 25-fold in male pericentral hepatocytes but only 4-fold periportally (**Fig. 6D**). The pericentral-specific expression of lnc4726 in female liver is thus recapitulated by cGH infusion in male liver, suggesting there is an intrinsic, underlying pericentral specificity of this lncRNA that is revealed when the gene is activated by exposure to GH in a persistent, female-like manner.

### TCPOBOP perturbs the sex-bias of many hepatic genes

TCPOBOP is a direct agonist of the nuclear receptor CAR [76], which shows pericentral-biased expression in male hepatocytes [60] and may contribute to the pericentral zonation of many xenobiotic metabolic reactions [75]. TCPOBOP induces widespread changes in liver gene expression within hours [45]. Here, we examined the impact of TCPOBOP exposure on sex-biased gene expression and zonation in mouse hepatocytes. CAR showed a pericentral bias in expression, both in control and in TCPOBOP-exposed hepatocytes, with the strongest zonation bias (4 to 5 fold) seen in male liver (**Fig. S6A**). Furthermore, CAR was expressed in a female-biased manner in periportal, but not pericentral hepatocytes, both with and without TCPOBOP exposure (**Fig. S6A**), which may contribute to the sex-dependent effects of TCPOBOP described below. RXRA, which forms a heterodimer with CAR, but also dimerizes with many other Nuclear Receptor family members, showed no sex differences in expression or zonation (**Fig. S6B**).

We compared the sex-biased transcriptomes of control versus TCPOBOP-exposed hepatocytes and identified many genes that gain sex-bias and many others whose basal sex-biased expression is lost following TCPOBOP exposure (**Fig. S7A**, **Table S5**). **Fig. 7** shows examples of genes that gain or lose sex bias, or whose sex bias is reversed in TCPOBOP-exposed hepatocytes. Rad51b gains female specificity due to its suppression by TCPOBOP in both periportal and pericentral male hepatocytes, whereas lncRNA 13508 gains female specificity due to its stronger induction in periportal female than periportal male hepatocytes (**Fig. 7A**). In contrast, Hsp90ab1 gains male specificity due to its increased expression in male but not female TCPOBOP-treated hepatocytes, whereas lnc9497 acquires male specificity in both periportal and pericentral hepatocytes due to strong suppression (9 to 17-fold) by TCPOBOP in female hepatocytes (**Fig. 7B**). Lnc33519 exemplifies loss of sex bias: it is highly expressed in male but not female hepatocytes under normal physiological conditions, but after TCPOBOP exposure is strongly induced in female but not male liver, thereby abolishing its male-biased expression (**Fig. 7C**). Sex bias reversal and sex bias loss are both exemplified by G6pc, which shows significant female bias in control liver (**Fig. 7D**). In male liver, TCPOBOP induces G6pc in periportal hepatocytes but in female liver TCPOBOP represses G6pc in both periportal and pericentral hepatocytes. Consequently, TCPOBOP abolishes the female bias of G6pc pericentrally and reverses its sex bias periportally. Overall, TCPOBOP reversed the sex bias of 4.2% of genes that show sex bias in control liver (**Table S5**; examples in **Fig. S7C**). Several of these genes make important contributions to hepatic carbohydrate, steroid and fatty acid metabolism, including G6pc [77], Zbtb7c [78], Gna14 [79] and Elovl5 [80], and dysregulation of their sex-dependent expression could contribute to the complex hepatic metabolic alterations associated with TCPOBOP exposure. Finally, Jazf1, a zinc finger repressor that protects from high fat diet-induced hepatic steatosis [81, 82], has an intrinsic underlying sex-bias that is increased by TCPOBOP exposure; thus, Jazf1 is lowly expressed with a 3-fold female bias in untreated liver, but following TCPOBOP exposure shows a strong increase, 10.8-fold, in female periportal hepatocytes (**Fig. S7B**). A lncRNA divergently transcribed from Jazf1, lnc5154, shows female-specific induction by TCPOBOP in both periportal and pericentral hepatocytes, but its maximal expression is low (**Table S5**, and FPKM ∼1 in TCPOBOP-induced female liver chromatin [35]).

**Fig. 7.**
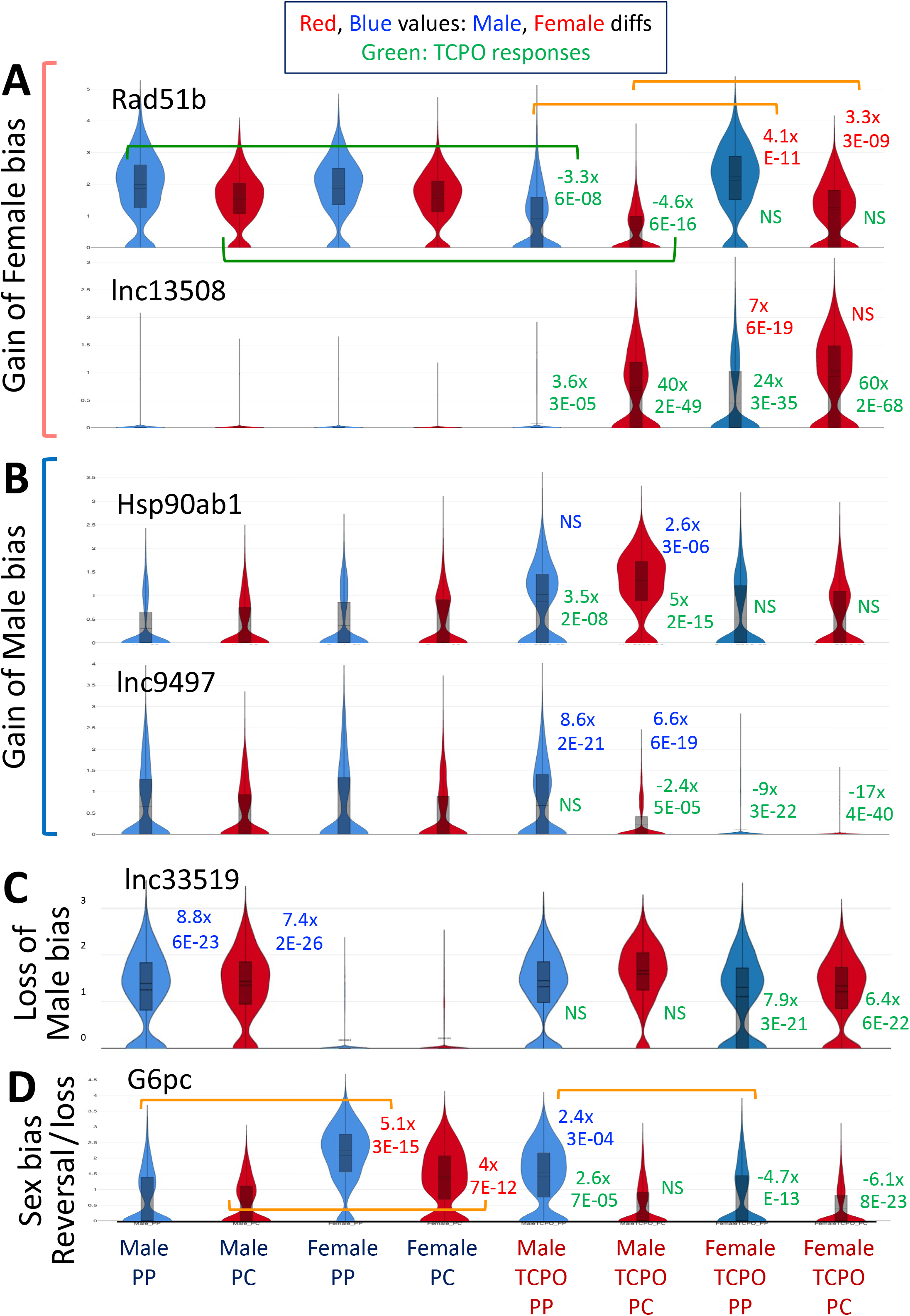
Genes that gain (A, B), lose (C) or reverse (D) sex-biased expression in either periportal (PP) or pericentral (PC) hepatocytes following TCPOBOP exposure. Shown to the right of the violin plots are fold-change and significance values for the sex bias in TCPOBOP-exposed liver (i.e., TCPOBOP-male versus TCPOBOP-female, evaluated separately for PP and PC hepatocytes) (genes in **A, B, D**) and the sex bias in control liver (gene in **C**), with red values indicating female bias and blue values male bias. Shown in green are fold-change and significance values comparing TCPOBOP-exposed to control, evaluated separately for PP and PC hepatocytes. The groups compared are marked by brackets for Rad51b and G6pc as examples. Positive green fold change values indicate gene induction by TCPOBOP, negative green fold change values indicate gene repression by TCPOBOP. NS, not significant.

### Impact of TCPOBOP on liver zonation patterns

We identified 1,613 hepatocyte zone-biased genes, including 828 zonated lncRNA genes, in TCPOBOP-treated liver – more than twice as many as the 719 genes showing zonation bias in vehicle (control) liver (**Fig. 8A**, **Table S3A**). More genes with a pericentral bias in control liver (n=142) lose their zonation bias in TCPOBOP-exposed hepatocytes than genes with a periportal bias in control liver (n=78) (**Fig. 8A**), consistent with the pericentral bias of CAR expression (**Fig. S6A**). Further, sex-specific zonation patterns were more prevalent in TCPOBOP-exposed liver than in control liver (**Fig. S4C, S4D** vs. **Fig. S4A, S4B**), consistent with the widespread and complex effects of TCPOBOP on sex-biased gene expression described above. These effects of TCPOBOP on zonation reflect several different mechanisms (**Fig. 8B-8F**), including the following: gain in pericentral zonation due to stronger induction by TCPOBOP of pericentral than periportal gene expression (Klhl33, where there is also a gain of female-bias; and Rarb); gain in periportal zonation when TCPOBOP specifically suppresses pericentral expression (Slc2a2); loss of pericentral zonation due to a preferential suppression of pericentral gene expression (Akr1c20); and loss of pericentral zonation due to much stronger induction of gene expression in periportal than pericentral hepatocytes (Cyp2c29; 115-fold vs. 6-fold increase in males). Thus, we conclude that TCPOBOP perturbs both the sex-specificity and zonation-bias of hepatocytes, most notably for pericentral-biased genes, and with major impact on many protein coding genes and lncRNAs.

**Fig. 8.**
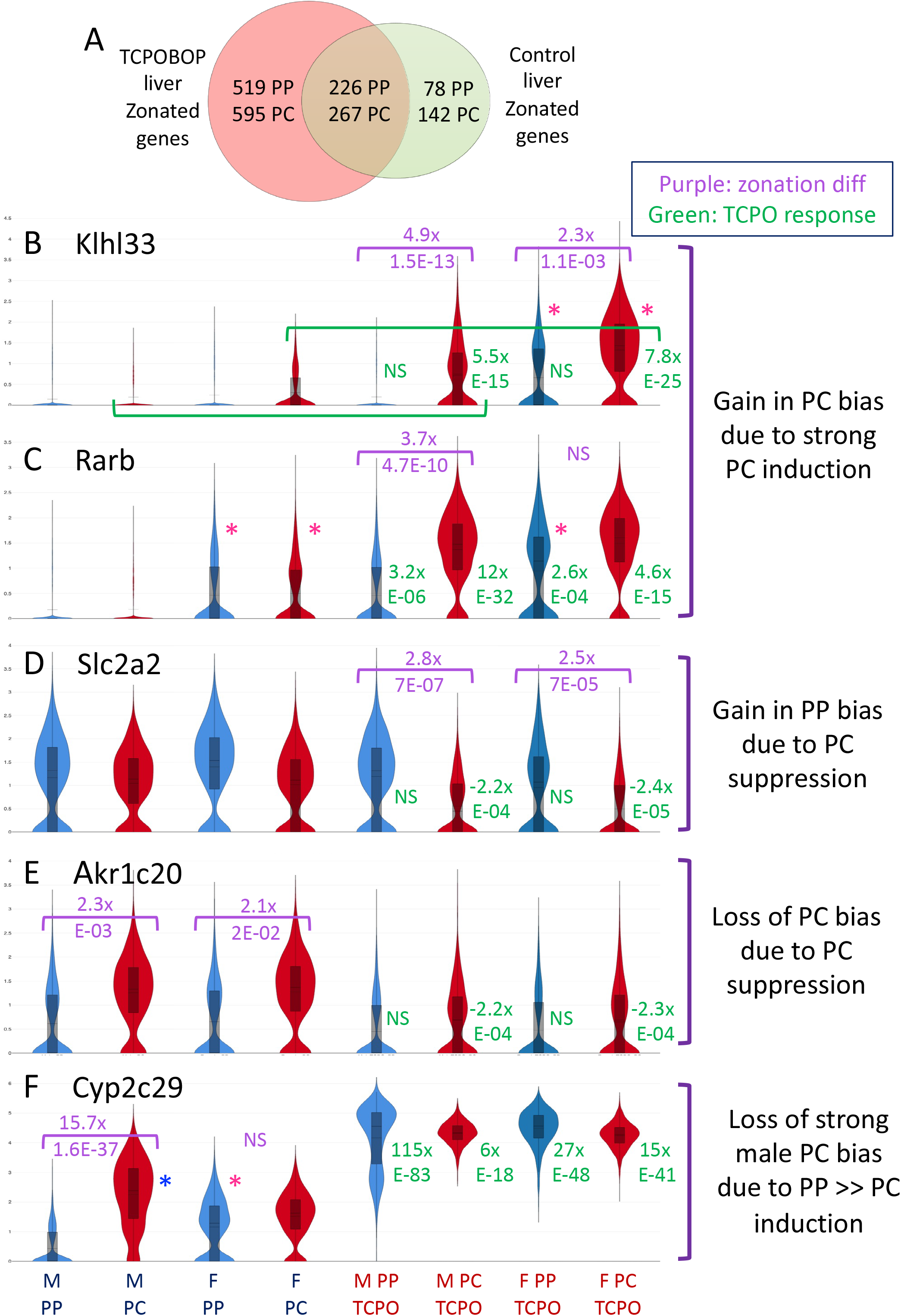
Changes in hepatocyte zonation in TCPOBOP-exposed liver. (**A**) Venn diagram presenting the number of zonation-biased genes identified in hepatocytes from either male or female TCPOBOP-treated liver and their overlap with zonation-biased genes identified in hepatocytes prepared from control liver (data from **Table S3A**, column I). Not included are 6 genes whose zonation patterns reverse following TCPOBOP exposure. Shown are examples of genes that show significant zonation-biased expression in TCPOBOP-treated liver but not in vehicle control liver (**B, C, D**), or vice versa (**E, F**), as indicated at the right. Purple values are fold-change and significance values for expression in periportal (PP) versus pericentral (PC) hepatocytes, evaluated separately for both control livers and TCPOBOP-exposed livers, in both male and female livers, as marked along the X-axis. Green values are fold-change and significance values comparing expression between TCPOBOP-exposed and control hepatocytes, evaluated separately for both periportal and pericentral hepatocytes, in both male and female livers, as marked in **B** for Klhl33 as an example. Significant sex difference in expression in either PP or PC hepatocytes, from either control or TCPOBOP-exposed liver, is marked by a pink asterisk (female-biased expression) or a blue asterisk (male-biased expression). NS, not significant.

### Impact of TCPOBOP on sex-biased hepatocyte function

Functional annotation clustering was used to identify key functional features of the sex-biased transcriptome in control liver and changes that occur following exposure to TCPOBOP (**Table S6A**). First, given the large number of new sex-biased genes revealed by the present snRNA-seq study, we compared the biological processes and molecular functions enriched in the sex-biased gene sets identified by snRNA-seq (**Table S2B**) to those identified by bulk liver RNA-seq (**Table S2D**). Very similar results were obtained for the two male-biased gene datasets; however, for the female-biased gene sets we found that snRNA-seq gave much stronger enrichment of lipid metabolic processes, including many genes active in cholesterol and fatty acid biosynthesis and metabolism (69 female-biased genes in annotation cluster, Enrichment score (ES) = 15.3; top term Bonferroni p-val = 6E-19) than did the female-biased bulk RNA-seq dataset (16 genes in annotation cluster, ES = 3.2; top term Bonferroni p-val = 0.03) (**Table S6B**). Many of the novel female-biased genes in this cluster have either a moderate to low sex bias (< 3-fold), low expression, or show sex-bias in only one hepatocyte cluster, and/or they have significant non-sex-biased expression in an NPC population, all of which tend to decrease the sensitivity for detection of sex-biased expression by bulk RNA-seq.

Many genes either gain or lose sex-biased expression following TCPOBOP exposure, as discussed above. Strikingly, the set of genes that lose female-bias is highly enriched in steroid biosynthesis and lipid metabolic process genes (ES = 15.2, top term Bonferroni p-val = 2.8E-20) (**Table S6E**), while the genes that gain female bias are weakly enriched for a cluster with 20 immune response genes (ES = 2.65; Bonferroni p-val < 0.05) (**Table S6F**). In contrast, the genes that gain male bias showed strong enrichment for a set of 13 stress response genes, including 8 heat shock protein genes (ES = 8.58, Bonferroni p-val = 3.63E-10) (**Table S6K**). These sex-differential effects of TCPOBOP likely contribute to sex differences in the effects of CAR activation on liver physiology and pathology.

Finally, we compared the genes that acquire zonation bias in TCPOBOP-treated males versus females. Functional annotations for genes that gain a periportal-biased expression pattern in male but not female hepatocytes include the terms biosynthesis of amino acids and glucagon signaling/insulin resistance (**Table S7A**), which are both associated with periportal hepatocytes in control liver [83], while the genes that acquire pericentral-biased expression in males but not females include the top term chemical carcinogenesis (**Table S7B**). Of note, the increased periportal bias of glucagon signaling/insulin resistance in TCPOBOP-exposed male hepatocytes reflects the preferential repression of these genes in pericentral hepatocytes (**Table S7C**). This effect is exemplified by Slc2a2 (**Fig. 8D**), a key liver glucose transporter, and by G6pc (**Fig. 7D**), a key sex-dependent enzyme of glucose homeostasis. Genes that acquire periportal-biased expression in female but not male hepatocytes from TCPOBOP-exposed liver show enrichment for the annotation term circadian rhythm (**Table S7D**). While the core clock genes controlling circadian rhythm do not show zonated expression in mouse hepatocytes [84], a result confirmed here in both male and female control liver, we did observe a gain of periportal zonation for 4 of 9 core clock genes following TCPOBOP exposure (**Table S7F**).

## Discussion

The metabolic zonation of hepatocyte function and its regulation by factors such as Wnt/β-catenin, hypoxia inducted transcription factors and hedgehog signaling are well established [75, 85, 86], but until recently, global analysis of hepatocyte zonation patterns of individual genes was not practical. The introduction of single cell sequencing technologies and associated methods for data analysis and integration [87] has led to major advances, including the discovery of large numbers of protein coding genes showing zonated expression in hepatocytes from male mice [60] and in studies using human liver tissue [88, 89]. Differences in zonation between males and females have been observed by laser capture microdissection [90] and may impact liver diseases, such as fatty liver disease, non-alcoholic steatohepatitis and hepatocellular carcinoma, which show significant sex differences in their incidence and severity, both in rodent models [11, 91, 92] and in humans [11, 93–95]; however, a comprehensive, single cell-based analysis of underlying sex differences is lacking. Here, we address this gap using single nucleus RNA-seq analysis to further our understanding of the interaction between liver zonation and sexually dimorphic gene expression, and how these complex, regulated expression patterns can be perturbed by xenobiotic exposure. We isolated nuclei from pools of frozen liver tissue for each of five biological conditions and utilized 10x Genomics technology to prepare snRNA-seq libraries and obtain high quality transcriptomic data for 32,000 nuclei with a mean of 2,000 genes detected per nucleus. Sequenced libraries were normalized, integrated across biological conditions using Harmony [96], and clustered to identify nine distinct liver cell-type-specific clusters, including four major hepatocyte clusters comprising 65% of all cells. Differential expression analysis between male and female cell clusters established that liver sex-biased gene expression is largely restricted to periportal and pericentral hepatocytes, where we found key liver transcription factors and other known regulators of sex-biased gene expression are most highly expressed. Moreover, many genes showing zonation of hepatocyte expression exhibited their zonation bias in only one sex. Infusion of male mice with GH for seven days, an established approach to feminize the sex-dependent plasma GH profiles, was effective in feminizing gene expression in both periportal and pericentral hepatocytes, with large numbers of female-biased genes induced and male-biased genes repressed. Finally, TCPOBOP, a foreign chemical agonist ligand of the Nuclear Receptor CAR (Nr1i3), induced widespread dysregulation of gene expression, including changes leading to both a gain and loss of sex-biased expression and zonation-bias for many genes, with loss of zonation more common for genes with a bias for pericentral hepatocytes, where CAR is more highly expressed.

Prior single cell-based studies of liver cell heterogeneity and zonation largely utilized single cell (sc) RNA-seq, rather than single nucleus (sn) RNA-seq [58, 60, 61, 88, 89]. Here, we implemented single nucleus sequencing with two specific goals in mind: first, to capture nuclei representing all cells present in bulk liver tissue, without introducing the bias with respect to recovery or viability of individual cell types that occurs when cells are isolated from dissociated liver tissue [97]; and second, to increase the sensitivity for single cell-based analysis of lncRNAs, a majority of which are strongly enriched in nuclei compared to cytoplasm [35]. The increased sensitivity of lncRNA detection gained by using snRNA-seq enabled us to discover and characterize many novel sex-biased, zonated, and TCPOBOP-responsive lncRNAs, which may serve as targets for future functional studies. While prior liver scRNA-seq studies have yielded cell type specificity data for certain highly expressed lncRNAs with established biological functions, we believe this is the first study to systematically examine on a global scale single cell-based lncRNA expression profiles in liver tissue for more than 48,000 liver-expressed lncRNAs. Remarkably, we detected 35,808 (75%) of these liver-expressed lncRNAs in one or more of the five biological conditions interrogated in the present snRNA-seq study. Notably, the number of zonation-biased lncRNA genes detected in our study was up to 12 times greater than we could identify from the Tabula Muris liver dataset [67], a widely used reference standard, in large part due to the use of nuclei rather than cells in our analysis.

One potential limitation of sequencing nuclei is the anticipated decrease in sensitivity to detect genes whose RNA transcripts are enriched in the cytoplasm as compared to the nucleus [35]. Indeed, we were unable to detect several of the Wnt signaling pathway factors that regulates liver zonation, as well as several liver-expressed transcription factors of interest. Top enriched key terms and functions for cytoplasm-enriched mRNAs include mitochondrial function, ribosomal proteins, and protein transport (Table-S2F of [35]). In contrast, single cell nuclear sequencing is expected to increase sensitivity for detection of genes that generate nuclear-enriched mRNAs, whose top enriched terms are glycoproteins, membrane, and extracellular matrix (Table-S2G of [35]).

We observed a higher ratio of pericentral to periportal hepatocytes in female than in male mouse liver. This sex difference, which was not previously described, could contribute to sex bias in expression of genes and biological pathways specifically associated with pericentral hepatocytes [56, 57] and to sex differences in susceptibility to chemical exposures with hepatocyte zone-dependent toxicities [50–52]. The relative decrease in pericentral hepatocyte abundance in males was seen in both control and TCPOBOP-treated livers, and when comparing untreated male livers to livers from male mice feminized by cGH infusion. Further, we observed the same sex difference when analyzing the Tabula Muris hepatocyte dataset [67], which is single-cell based, rather than nucleus-based. In the latter dataset, the higher pericentral to periportal hepatocyte ratio seen in female liver is apparently associated with a shift in hepatocyte number in female liver from the mid-lobular to the pericentral region. Hepatocyte zonation is a continuum that stretches from the periportal vein to the central vein [85], and accordingly, we found that the pericentral cell population that bordered the periportal region displayed more mid-lobular characteristics. Moreover, the female bias in pericentral cell number was primarily associated with increased cell numbers in the pericentral cell sub-cluster closest to the periportal cluster, which has more mid-lobular characteristics (**Table S1C**). Another factor in the increased representation of pericentral hepatocytes in female liver seen in our dataset could be the presence of bi-nucleated hepatocytes [85, 98], which are more prevalent in female liver [64, 65], and would consequently increase pericentral nuclei counts in females.

We identified two hepatocyte-related cell clusters of special note. One cluster was enriched for expression of the DNA topoisomerase Top2a [68], a characteristic marker of dividing cells, and showed a 40-50% increase in abundance in livers from mice treated with TCPOBOP, which stimulates liver cell proliferation [69, 70]. We identified this cluster as dividing hepatocytes based on its enrichment for hepatocyte marker genes, primarily those from pericentral and mid-lobular hepatocytes. This finding is consistent with the preferential proliferation of periportal hepatocytes that occurs following pericentral hepatocyte damage, which may be occurring here in TCPOBOP-exposed liver [99]. A second hepatocyte-related cluster was characterized by strong enrichment for hepatocyte secretory proteins but low to moderate levels of liver transcription factors and other hepatocyte marker genes [60]. This cluster appears to represent nuclei non-specifically associated with membrane-bound polyribosomes and their tethered mRNAs encoding membrane-bound and secretory proteins.

Sex-biased gene expression was restricted to hepatocytes, as seen for 93% of 1787 sex-biased genes. This cell type specificity may in part be explained by the elevated expression in hepatocytes of key sex-specific or GH-regulated transcription factors, as well as several liver-enriched transcription factors implicated in sex-biased gene expression (**Fig. S2**). Many sex-biased genes showed zonation-biased expression within the liver lobule and constituted almost half of genes showing zonation-biased expression, highlighting the interconnection of these two mechanisms of gene regulation. Further, many sex-biased genes showed zonation-biased expression in only one sex. One striking example, Cyp2c29, showed strong pericentral-biased expression in male liver, but no significant zonation in female liver; this resulted Cyp2c29 having a male-biased expression pattern in pericentral hepatocytes but female-biased expression in periportal hepatocytes by a mechanism that most likely involves a combination of zonation and sex-biased regulators. Consistent with this, in cGH-infused male liver, Cyp2c29 was selectively induced in periportal hepatocytes, abolishing the zonation bias and recapitulating the expression pattern of female liver. Overall, cGH treatment down regulated a substantial subset of male-biased genes while inducing the expression of many female-biased genes, recapitulating our earlier finding by bulk liver RNA-seq analysis [31].

In a prior study examining sexual dimorphism of liver zonation via laser capture microdissection, only the periportal region was found to show sex differences in metabolic functions [90]. Two upstream regulators of zone-specific genes were found to be sexually dimorphic, namely, the periportal-expressed and male-biased Srebf1, and the pericentral, female-biased transcription factor Trim24 [90]. Here, we found a 2-3 fold female-bias in the expression of Trim24 in both pericentral (higher expression) and periportal hepatocytes, but did not observe any zonation-bias or sex-bias for Srebf1.

TCPOBOP induced widespread changes to the sex-biased transcriptome, with many genes showing a gain or a loss of sex-biased expression, as well as changes in zonation. Notably, genes that lost female-biased expression, but not genes that lost male-biased expression, following TCPOBOP exposure were highly enriched (15-fold) for steroid biosynthesis and lipid metabolism differences, which may contribute to the sex-dependent effects of TCPOBOP exposure. Quantitative differences in response between the sexes can be anticipated from the higher levels of CAR protein reported for female compared to male mouse liver [100, 101], but the widespread sex-dependent effects of TCPOBOP seen here indicate other, more complex mechanisms are at play, consistent with our earlier findings using bulk liver RNA-seq [45]. Nearly twice as many pericentral-biased genes as periportal-biased genes showed a loss of zonation-bias following TCPOBOP exposure in either male or female liver, consistent with the strong, 4.3-fold pericentral zonation that we found for CAR, but not its dimerization partner RXRA, in male liver. However, for many genes, TCPOBOP induced large changes in expression in both periportal and pericentral hepatocytes, a subset of which were sex-dependent and GH-regulated. Other chemical exposures preferentially affect pericentral hepatocytes and thereby dysregulate liver zonation, such as pericentral liver fibrosis induced by exposure to carbon tetrachloride, a known hepatotoxicant [102], in that case attributed to the pericentral expression of Cyp2e1 (confirmed here: 5.2-8.5-fold,FDR < E-12; **Table S3**) and its activation of carbon tetrachloride to reactive free radical metabolites [103]. Similarly, chronic exposure to 2,3,7,8-tetrachlorodibenzo-p-dioxin (TCDD) led to large changes in gene expression in many liver cell types, with changes to hepatocyte zonation [104].

One limitation of this study is the sensitivity for detection of RNAs expressed at a low level, including many lncRNAs, which remains a challenge, despite the increased sensitivity achieved by using snRNA-seq.

Nevertheless, we were able to analyze and characterize many such lowly expressed lncRNAs. One example is lnc-LFAR1, which is functional in promoting liver fibrosis [66, 105] and based on our analysis was detected in fewer than 10% of hepatocytes (log normalized signal intensity 0.09 UMI/cell cluster). Feature plot analysis revealed that this liver lncRNA is expressed in a male-specific manner in both control and TCPOBOP-exposed liver and is repressed to female levels by cGH infusion, confirming its hormone-regulated sex bias; moreover, these patterns were validated and shown be robust by statistical analysis (**Fig. S8**). Conceivably, this lncRNA may contribute to the male-bias in diseases such as non-alcoholic steatohepatitis and the development of liver fibrosis [11].

In conclusion, by utilizing nuclei, we were able to discover hundreds of lncRNAs with unique sex-biased and zonation-biased expression patterns, many of which responded to hormone-based feminization of the liver by cGH infusion or responded to TCPOBOP exposure. Overall, our findings highlight the interconnectedness of sex-bias and zonation-bias in the mouse liver and exemplify how these natural patterns of expression respond to endogenous hormones and can be dysregulated by exposure to foreign chemicals.

## Methods

### Animal experiments

All mouse work was carried out in compliance with procedures approved by the Boston University Institutional Animal Care and Use Committee (protocol # PROTO201800698), and in compliance with ARRIVE 2.0 Essential 10 guidelines [106], including study design, sample size, randomization, experimental animals and procedures, and statistical methods. CD-1 mice between 7-9 wk of age (strain Crl:CD1(ICR)) were purchased from Charles River Laboratories (Wilmington, Massachusetts). Frozen livers collected by Dr. Hong Ma of this laboratory were from male and female mice, either vehicle-treated or exposed to TCPOBOP. TCPOBOP (1,4-bis(2-(3,5-dichloropyridyloxy))benzene) (Santa Cruz Biotechnology; Chem Cruze, Cat. #SC-203291) was dissolved in DMSO at 7.5 mg/ml and then diluted 10-fold into corn oil, followed by IP injection at 4 µl per gram body weights (final dose: 3 mg TCPOBOP and 4 µl of 10% DMSO in corn oil, per kg body weight). Injections were performed at 8 AM and mice were euthanized 27 h later, at 11 AM (Boston University animal facility light cycle: 7:30 AM to 7:30 PM). Livers from an independent set of male CD-1 mice, untreated males and 7-day cGH-treated males, were those described previously [107]. Mice were implanted with a subcutaneous Alzet model 1007D osmotic mini-pump set to deliver 20 ng of recombinant rat GH per h per gram body weight for 7 d [22, 107]. Mice were euthanized, livers were collected and flash frozen in liquid nitrogen then stored at -80°C.

### Isolation of intact single nuclei for sequencing

Single nuclei were obtained from flash frozen mouse liver tissue based on a protocol for isolating nuclei from frozen kidney tissue [108]. The protocol was optimized for mouse liver, based on a method to isolate nuclei from mouse brain tissue provided by Christine Cheng (Dept. of Biology, Boston University). The following buffers were prepared fresh daily in advance and kept on ice for up to 2 h: **Base Solution,** 10 mM Tris-Cl, pH 7.4, 146 mM NaCl, 1 mM CaCl_2_ 21 mM MgCl_2_; **Lysis Buffer,** Base Solution containing 0.1% Triton X-100 (Cat. #T8787, Sigma), with 80 U/mL Protector RNase Inhibitor (Cat # 3335402001, Roche) added just prior to use; **BSA Wash Buffer,** 1X PBS containing 2% BSA and 0.02% Tween- 20, with 80 U/mL Protector RNase Inhibitor added just prior to use. We prepared 6 single nucleus sequencing libraries, one for each of the 6 treatment groups studied: control (vehicle-treated) male liver, 27-h TCPOBOP-exposed male liver, control (vehicle-treated) female liver, 27-h TCPOBOP-exposed female liver; and prepared in a separate batch: untreated male liver (control group, marked Male1 in the figures) and 7-day cGH-infused male liver. Nuclei were prepared for each treatment group from frozen liver tissue collected and pooled from n = 4 biological replicate livers (50-75 mg frozen liver tissue from each of 4 mice per treatment group). Liver samples were cut and stored in an Eppendorf tube on dry ice until ready for extraction of nuclei, to minimize premature thawing and RNA degradation. Each of our snRNA-seq libraries thus represents snRNA-seq expression profiles from livers derived from a population of four mice, rather than a single individual, minimizing the impact of individual variation on the single cell and differential expression transcriptomic profiles and datasets obtained. All pipette tips used were RNase-free and DNase-free.

The 4 frozen pre-cut pieces of liver comprising each group were transferred to a 3 mL glass-on-glass Dounce homogenizer with 1 ml of Lysis Buffer, on ice. Keeping the homogenizer on ice, the samples were dounced for 10 strokes with pestle A (loose fit) followed by 10 strokes with pestle B (tight fit) or until fully homogenized. BSA Wash Buffer (1 ml) was then added, the sample then pipetted up and down a few times to mix, and then passed through a 40 µm cell strainer into a 50 ml conical tube on ice. Three additional washes of BSA Wash Buffer (1 ml) were used to rinse the homogenizer and pestles on ice and then passed through the same 40 µm cell strainer and added to the homogenized sample in the conical tube. The homogenizer and pestles were washed with MilliQ water on ice three times before processing the next sample.

Homogenized samples were kept on ice until all samples were ready to proceed to the next stage. Each homogenized sample was divided into three 1.5 ml Eppendorf tubes, which were centrifuged at 500g in a swinging bucket centrifuge for 5 min at 4° C to pellet the lysed cells. The supernatant was discarded and pelleted nuclei then resuspended in 1 mL BSA Wash Buffer, pooling all three tubes back into a single sample. Half of the sample, 500 µL, was transferred to a new 1.5 mL Eppendorf tube then diluted with 500 µL BSA Wash Buffer before pelleting at 500g for 5 min at 4° C. The supernatant was discarded and the pellet was resuspended in 1 mL BSA Wash Buffer then passed through a 20 µm cell strainer on ice (Cat # 43-50020-03, PluriSelect). The strained sample was transferred to a LoBind 1.5 mL Eppendorf tube (Cat # 022431021, Eppendorf) on ice, visualized (**Fig. S1A**), quantified using a Cellometer (Nexcelom) and diluted to a final concentration of 1.2 x 10^6^ nuclei/mL in BSA Wash Buffer. Samples were kept on ice up to 1 h and transferred to the Boston University Single Cell Sequencing Core for 10X Genomics single nucleus sequencing.

### Single nucleus sequencing (snRNA-seq)

Nuclei diluted to approximately 1.2 x 10^6^ nuclei/mL were recounted using a hemocytometer and loaded onto a 10x Genomics Chromium instrument to prepare Single Cell 3’ Sequencing (version 3 chemistry) libraries (**Table S1A**). A Bioanalyzer was used to verify concentration and size distributions prior to Ilumina sequencing at Novogene, Inc (Sacramento, CA). Sequencing libraries from male and female livers (vehicle control or TCPOBOP exposed) were processed and sequenced separately a second set of samples, comprised of nuclei from untreated control male livers (designated ‘Male1’) and cGH-infused male livers. Raw sequencing files, along with processed data in the form of a 10X Genomics Cloupe file (see below) are accessible at GEO (https://www.ncbi.nlm.nih.gov/geo/) and will be released upon acceptance for publication, accession # GSE188488.

### snRNA-seq data processing

Raw fastq sequencing files were processed using CellRanger (v3.1.0) and the Cellranger mkref pipeline to align sequence reads to mouse reference genome mm10. Gene annotations were input using a custom reference Gene Transfer Format (GTF) file containing 76,011 mouse genes. This GTF file is comprised of mouse mm10 gene coordinates for RefSeq protein coding genes (n = 20,960), mitochondrial genes (n = 13), a set of mouse liver-expressed lncRNAs genes (n = 48,261) that is more complete than the set described previously [45], RefSeq non-coding genes (NR accession numbers) that do not overlap with the set of 48,261 lncRNAs (n = 2,077), Ensembl non-coding lncRNAs that do not overlap either the RefSeq NR gene set or the 48,261 lncRNA gene set (n =4,697), 3 other lncRNAs with interesting liver functions (lnc-LFAR1, LeXis, Lnclgr), and 91 ERCC control sequences [109]. Two different GTF files comprising these 76,011 genes were used in our analyses (see Supplemental files).

GTFA_Exon_MouseLiver_scRNAseq.gtf, comprised of gene coordinates from all exonic regions in the set of 76,011 mouse genes, was used when analyzing the public Tabula Muris scRNA-seq data set [67] to count reads aligned to gene exons. GTFB_FullGeneBody_MouseLiver_snRNAseq.gtf, comprised of gene coordinates for the full gene body, including all exonic and all intronic regions of each gene in the set of 76,011 mouse genes, was used when analyzing our snRNA-seq datasets, to enable Cellranger to capture the many unspliced RNAs found in nuclei, including those derived from the 48,261 liver-expressed lncRNAs, in addition to partially processed and mature RNAs, many of which are abundant in the nucleoplasm [35]. As Cellranger excludes reads from genomic regions that overlap each other on the same strand, we made the following change for 4,352 overlapping genes that we identified (**Table S1F**), in order to minimize the impact of the exclusion of overlap regions from Cellranger counting. For these 4,352 genes, rather than using the full gene body for counting, which would lead to exclusion from two (or more) overlapping genes of sequence reads mapping to any region that is intronic to one gene but is exonic to the other, overlapping gene, we used a ‘modified gene body’ approach for counting, comprised of all exons of each gene plus all non-exonic regions that were unique to each gene. In this manner, sequence reads that map to an exon of one gene but overlap an intronic region of another gene are counted, as the overlapping intronic region of the second gene is excluded from the GTF annotation file for the second gene (**Table S1F**).

CellRanger statistics for all six snRNA-seq samples (**Table S1A**) indicate median values of 2,047 genes detected per nucleus, 45,560 total genes detected per biological condition, and 3,841 median UMI counts per nucleus. Sequencing data for all six samples was aggregated using CellRanger aggr, which enabled batch effect correction to be performed on the combined dataset. A normalized integrated count matrix was generated by dividing the unique molecular identifier (UMI) count for each gene by the total number of UMIs in each cell, followed by log-transformation. To fully integrate the datasets, Harmony [96] was applied to further remove unwanted variance by embedding and integrating shared cell types that are present across all six samples, by using the normalized integrated count matrix. Graph-based clustering of nuclei based on their gene expression profiles was performed using the FindClusters function in Seurat [110]. Uniformed Manifold Approximation and Projection (UMAP) plots (resolution 0.25, principal components = 10) were employed to visualize the clusters, and cell identities were assigned to each cluster using established marker genes (**Fig. 1**). Nuclei with >5% of UMIs mapping to mitochondrial genes were deemed lower in quality (e.g., due to mitochondrial contamination associated with apoptotic cell death or tissue necrosis) (n= 6,059 nuclei across the six snRNA-seq samples analyzed) and were removed. Additionally, a small cluster comprised of 1,316 nuclei across the 6 samples with higher expression of mitochondrial genes than the other remaining clusters was removed. DoubletFinder [111] was used to identify 2,563 nuclei likely to be comprised of at least two nuclei each (multiplets), which were removed (analysis parameters: top 10 principal components, 5% doublet formation rate). Together, these steps filtered out a total of 9,938 nuclei from an initial total of 42,131 nuclei across the six samples, leaving 32,193 nuclei for our analysis. These remaining nuclei were re-integrated using Harmony, and the UMAP then generated was uploaded to 10x Genomics Loupe Browser software (v 4.2.0) to produce a Cloupe file. This file, available at GEO (accession # GSE188488), was used for downstream analysis with the Loupe Browser based on 31,896 of the 32,193 nuclei, i.e., nuclei that mapped to the nine cell type clusters identified in **Fig. 1A**. Splitting of the aggregated UMAP by sample verified the effectiveness of Harmony-based integration, as indicated by the consistency of UMAP patterns for each cell type-based cluster for each of the 6 samples (**Fig. S1B**). Expression of the 13 mitochondrial genes in the final set of nuclei was visualized by dot blot analysis (**Fig. S1C**).

### Gene expression visualization and differential expression analysis

The Loupe Browser was used to project UMAPs of the cell type clusters obtained after integration and harmonization of the six samples. Expression levels of individual genes or small groups of genes (log normalized UMI counts) were visualized by dot plots, generated using the DotPlot function in Seurat [112], with dot intensity indicating expression level and dot diameter indicating the fraction of nuclei in each cluster that express the gene. Violin plots output from the Loupe Browser were used to display gene expression as the distribution of UMIs across all nuclei in each cell cluster after log normalization to all nuclei in the dataset, including nuclei in clusters not being compared in the figures. Values shown in black above each violin plot indicate the mean expression level associated with these globally log normalized violin plots; accordingly, given the inclusion of nuclei from other clusters in the normalization step, one cannot calculate differential expression between two clusters directly from the ratio of their mean cluster expression values. Rather, differential expression analysis between any two specific clusters was calculated using the Loupe Browser’s integrated Locally Distinguishing function. This method implements the negative binomial test based on the sSeq method [113] with Benjamini-Hochberg correction for multiple tests, and takes into account log-normalized average expression values as well as the distribution of UMIs across the two specific cell populations being compared to obtain fold-change and false discovery rate (FDR) values for each pair-wise comparison between cell clusters. Comparisons were made between clusters within a given sample (e.g., periportal vs pericentral hepatocytes for control male liver) and between samples (e.g., male periportal hepatocytes vs female periportal hepatocytes). Differential expression files output from Loupe Browser were processed using an in-house script and uploaded to an in-house gene expression database, named SEGEX, that was used to output the differential expression comparison experiments, including the Total Flag Sum (TFS) calculations capturing the expression patterns of each gene. These are presented in **Tables S2-S5** and explained in the legend to **Table S2B**. Results of these differential expression analyses for select genes are displayed in colored text to the right of the violin plots shown in each figure, except as noted. We did not apply the Remove Low Average Count gene filter option offered by the Loupe Browser to remove genes with average expression < 1 UMI per cell across each cluster, as we found it much too aggressive; namely, it eliminated many well-established sex-specific genes from our differential expression analyses. One limitation of this approach is that some of the lowest expressed genes identified as differentially expressed could be expressed at too low a level to be biologically relevant.

Differential expression between two cell clusters was used to identify genes showing sex-biased expression (**Table S2**), zonation-biased expression (**Table S3**), responses to cGH infusion (**Table S4**) and to characterize sex-biased and zonation-biased differential expression in TCPOBOP-treated male versus TCPOBOP-treated female hepatocytes (**Table S5**), all using an FDR filter of < 5%. In **Table S2E**, differential expression between the combined periportal and pericentral clusters (group 1) versus the 5 non-parenchymal cell (NPC) clusters (group 2) was performed using Loupe Browser to identify hepatocyte-specific genes and NPC-specific genes. To determine the total number of genes expressed at any level in our dataset, we performed global differential expression analysis between each cluster across all 6 samples (Globally Distinguishing feature comparison option within Loupe Browser); overall, 35,808 lncRNAs from the complete list of 48,261 liver lncRNAs (and 24,147 protein coding genes and other non-coding genes) were detected in one or more of the six snRNA-seq samples in our data set.

The Tabula Muris mouse liver scRNA-seq data set [67] was processed in the same manner as our mouse liver snRNA-seq dataset. Clusters were identified using the same list of cell-type specific markers shown in **Fig. 1C**, and differential expression was performed as described above to allow for comparison of the Tabula Muris data set to our data set.

### Functional annotation analysis

DAVID Bioinformatics Resources v6.8 Functional Annotation Tool was implemented using default parameters, including medium classification stringency. Output presented in the Supplemental tables includes all Annotation Clusters with an overall cluster Enrichment Score > 1.3, representing an FDR of < 0.05.

### Statistical analysis

Statistical methods for analysis of snRNA-seq data, including differential expression analysis between cell clusters within a given sample and between samples, as well as violin plot presentations and functional annotation analysis are described above.

## Abbreviations

CAR: constitutive androstane receptor, Nr1i3
FDR: false discovery rate
GH: growth hormone
cGH: continuous infusion of GH
lncRNA: long non-coding RNA
NPC: non-parenchymal cell
scRNA-seq: single cell RNA sequencing
snRNA-seq: single nucleus RNA sequencing
TCPOBOP: 1,4-bis(2-(3,5- dichloropyridyloxy))benzene
UMI: unique molecular identifier
UMAP: Uniformed Manifold Approximation and Projection

## Supplemental Figures

**Fig. S1.**
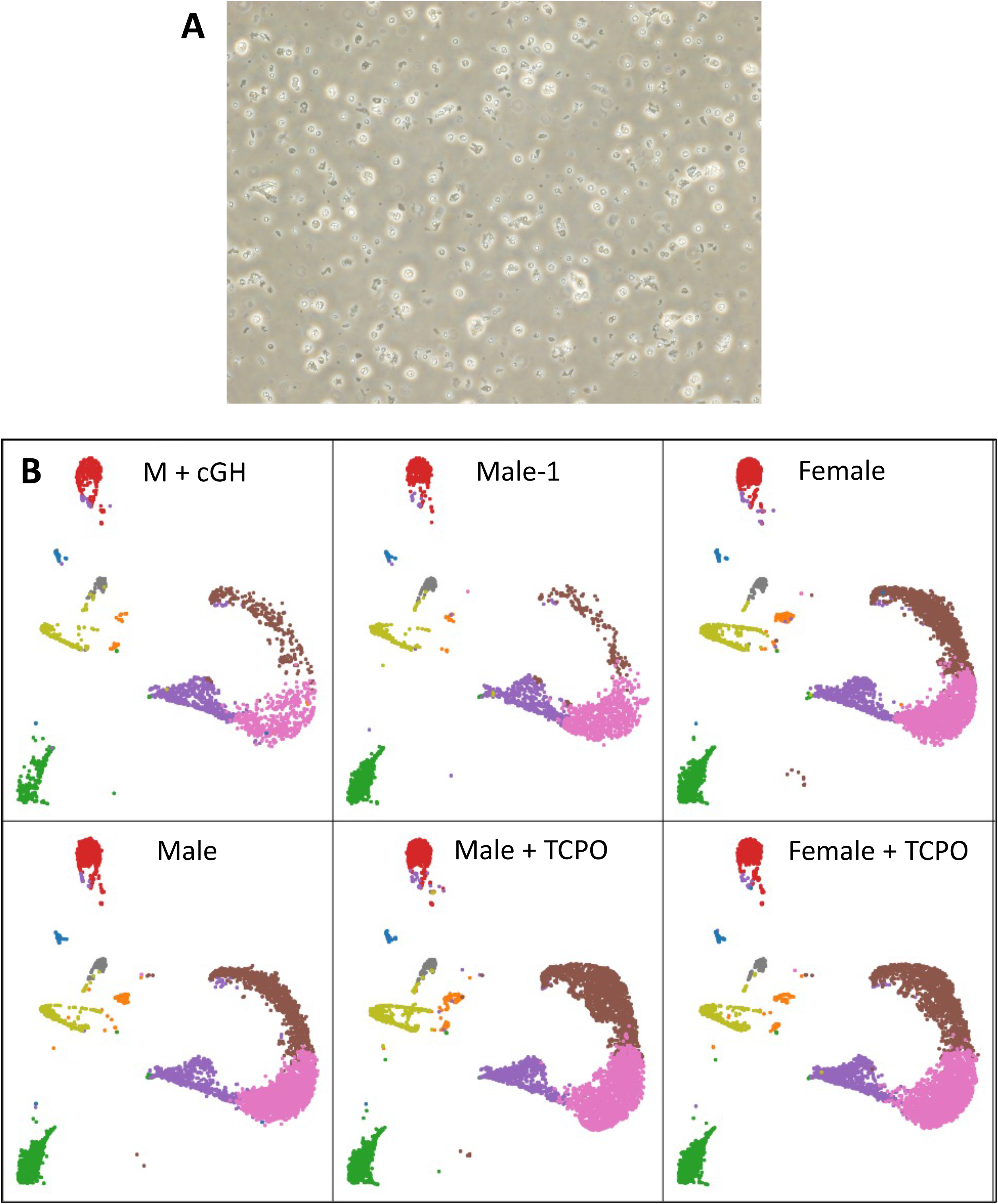
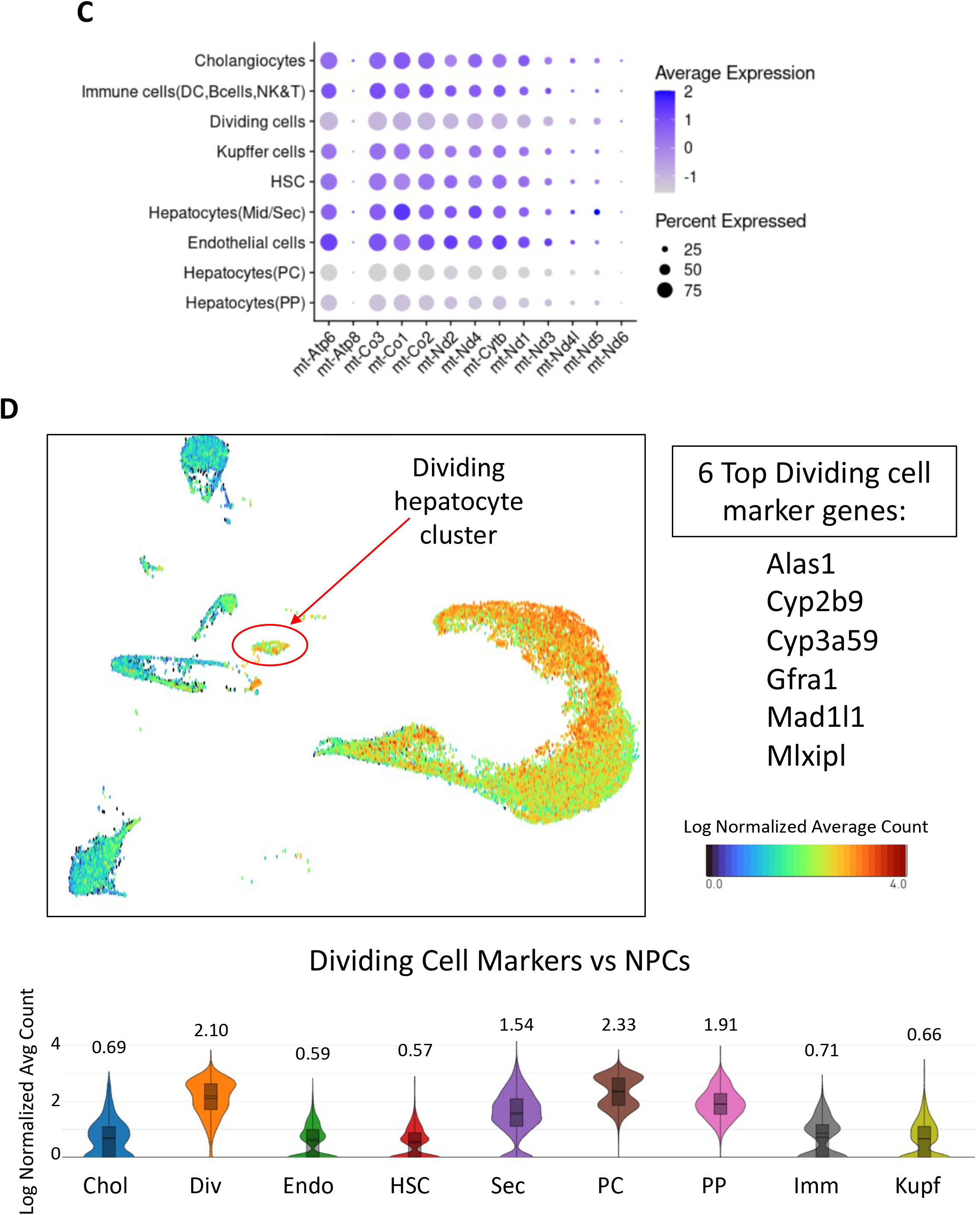
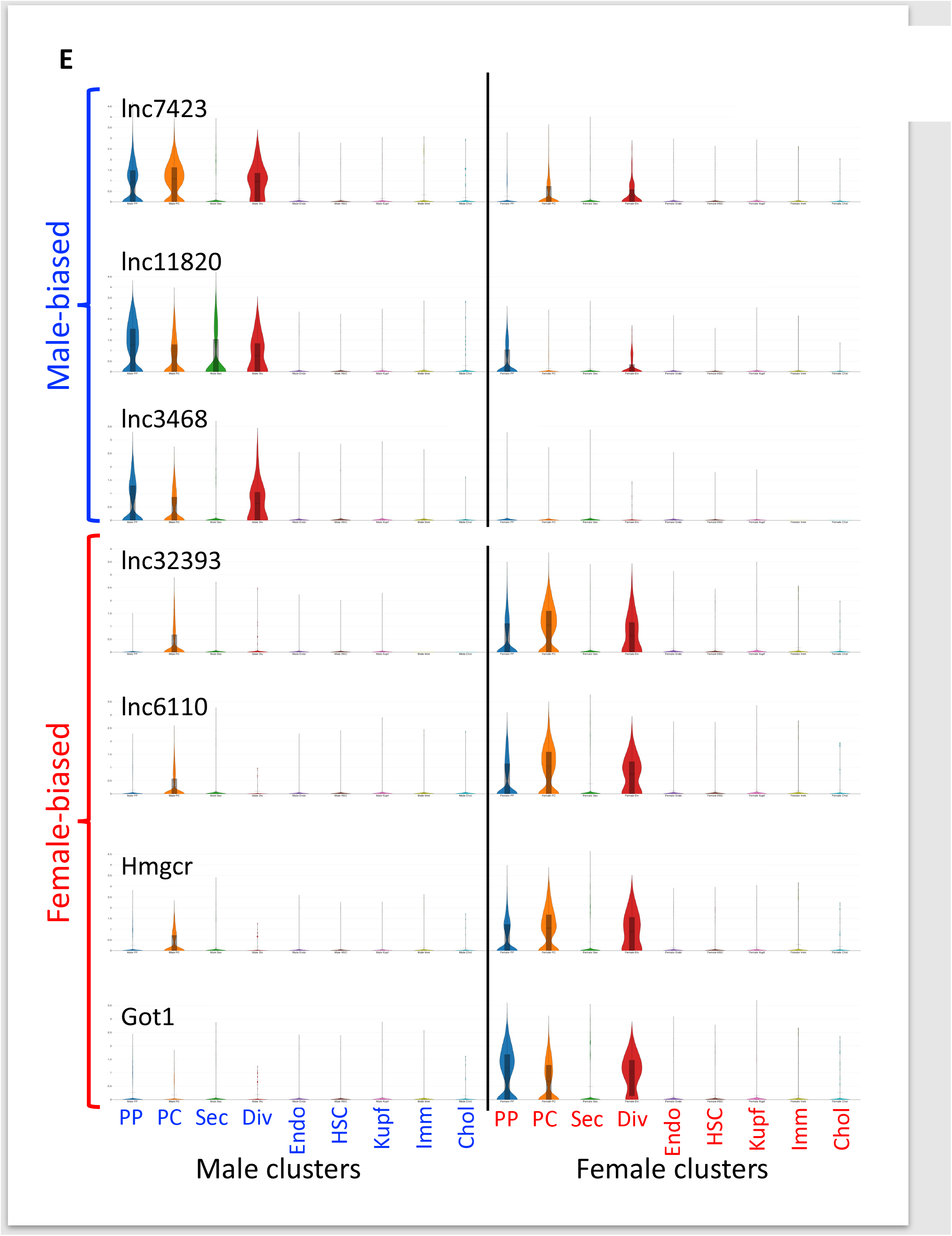
Isolated liver nuclei, mitochondrial content and dividing cell cluster. (**A**) Representative image showing the quality of nuclei isolated from frozen mouse liver tissue and then processed for snRNA-seq on the 10x Genomics platform. (**B**) Aggregated UMAP for the 6 snRNA-seq samples generated by Harmony (as in **Fig. 1A**) was split by sample to verify the effectiveness of Harmony-based integration, as indicated by the consistency of UMAP patterns for each cell type-based cluster for each of the samples. Legend identifies each cluster, with the number of nuclei shown in parenthesis. (**C**) Mitochondrial content in each of the 9 single nuclei UMAP clusters after removal of nuclei with >5% mitochondrial content. Dot plot shows the percentage of nuclei in each cluster that express each of the 13 mitochondrial genes (mt) indicated at the bottom (size of each spot) and the average expression level of each gene (color of each spot). (**D**) Cumulative expression level of 6 genes that are enriched in the dividing cells cluster (Div) when compared to the population of cells in the 5 NPC clusters: Endothelial (Endo), Hepatic Stellate Cells (HSC), Kupffer cells (Kupf), Immune cells (Imm) and Cholangiocytes (Chol). Expression data for all genes is shown in **Table S1E**. The UMAP indicates average log normalized expression of the combination of the 6 dividing cell marker genes in each nucleus, as per the color bar legend, evidencing that these markers are primarily expressed in hepatocytes, not NPCs. Violin plots at the bottom confirming this finding show log normalized average expression profiles for the set of 6 genes based on UMI counts per nucleus for all nuclei in the cluster. The average expression level in each cluster is indicated above each violin plot. (**E**) Examples of female-biased and male-biased genes whose expression is largely restricted to 3 hepatocyte clusters (PP, periportal; PC, pericentral; and Dividing cells). Also see **Fig. 2B-2C**.

**Fig. S2.**
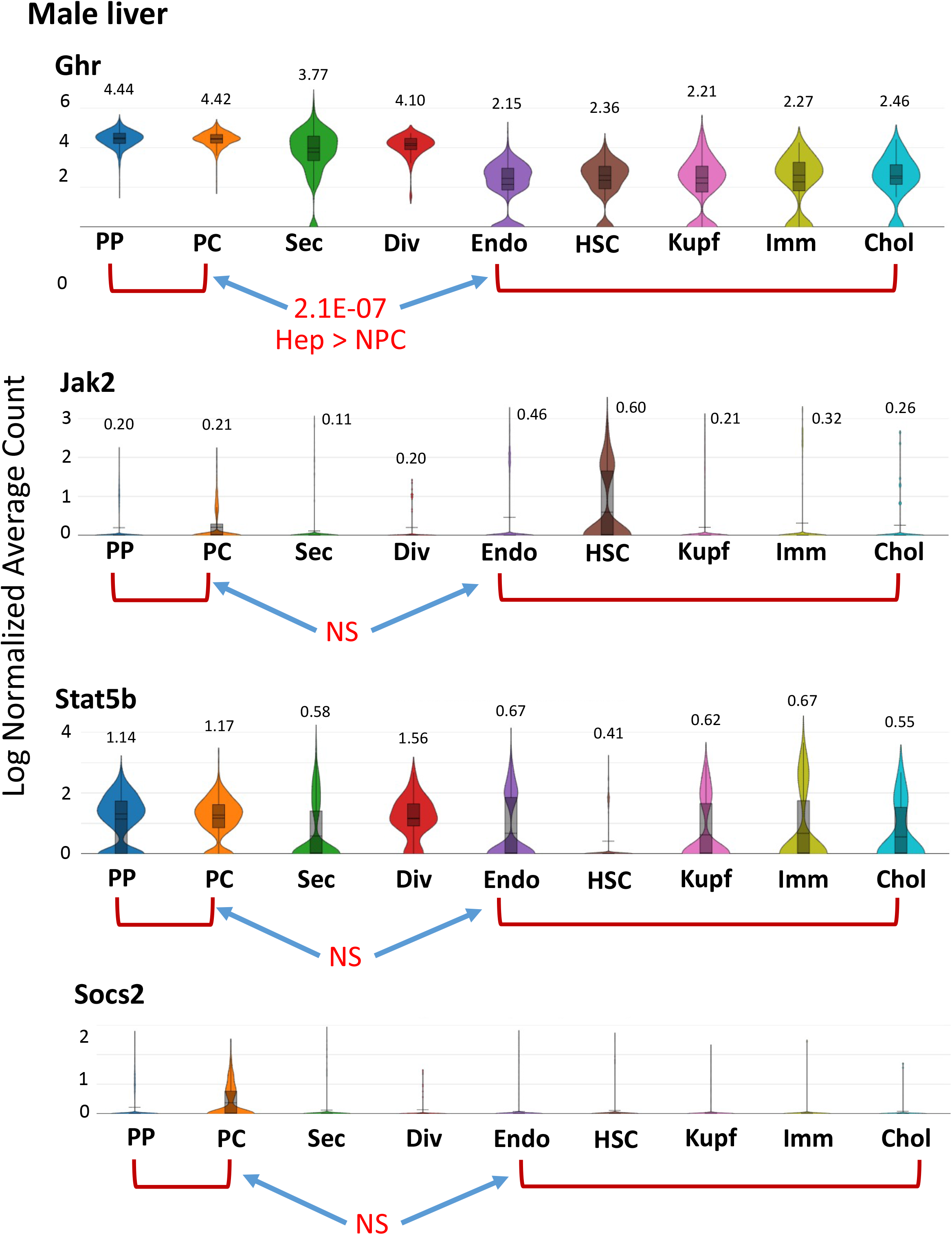
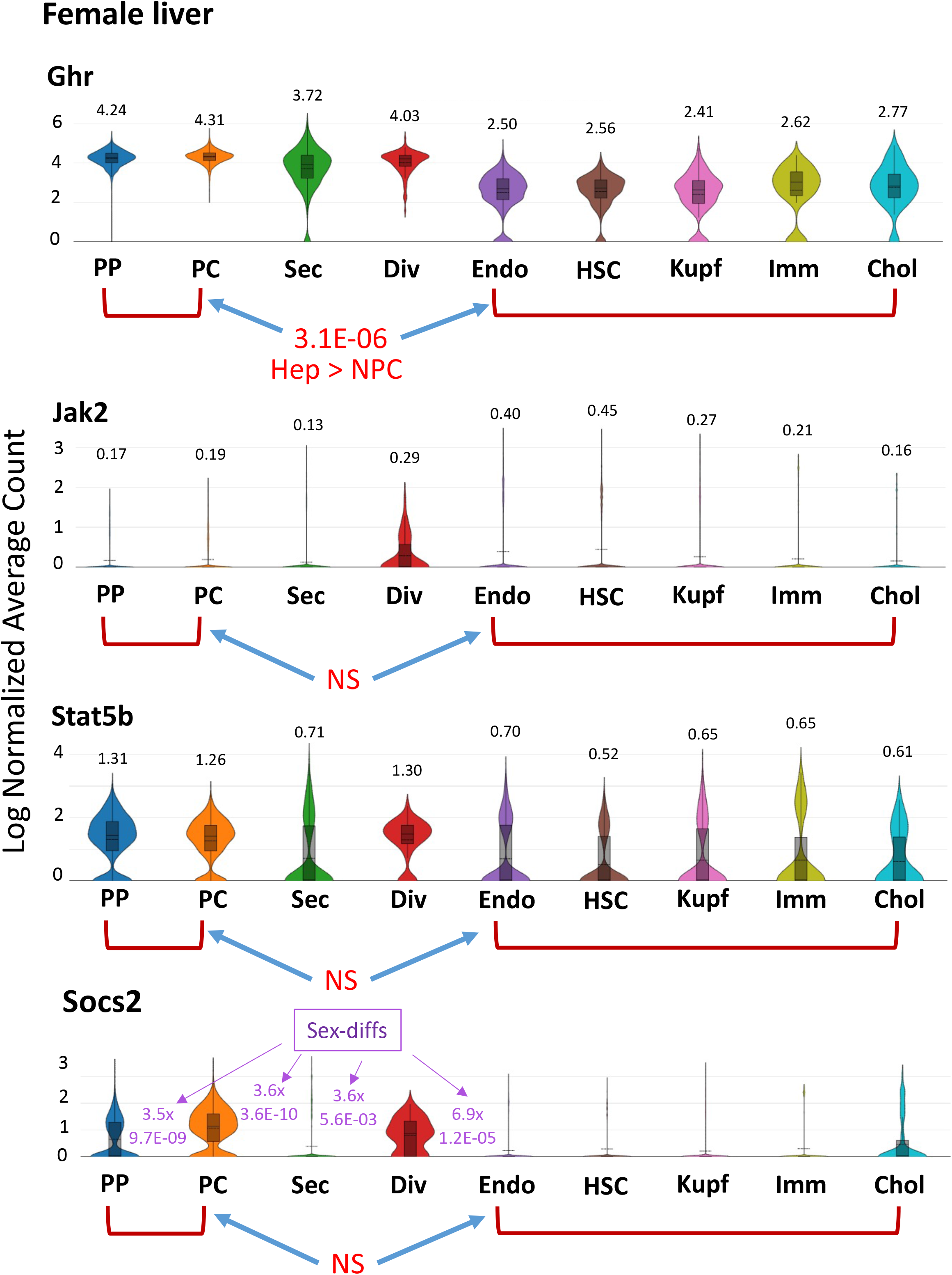
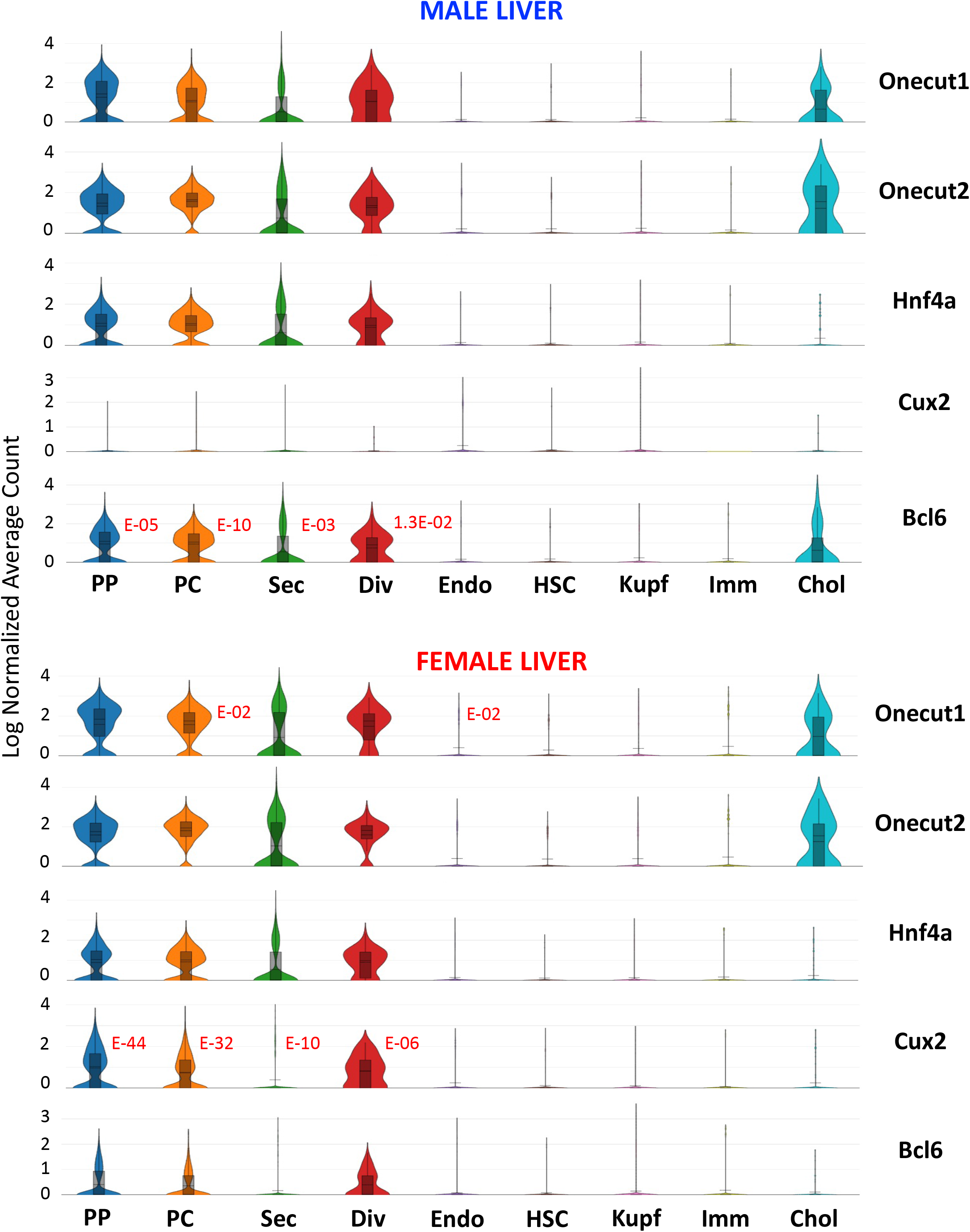
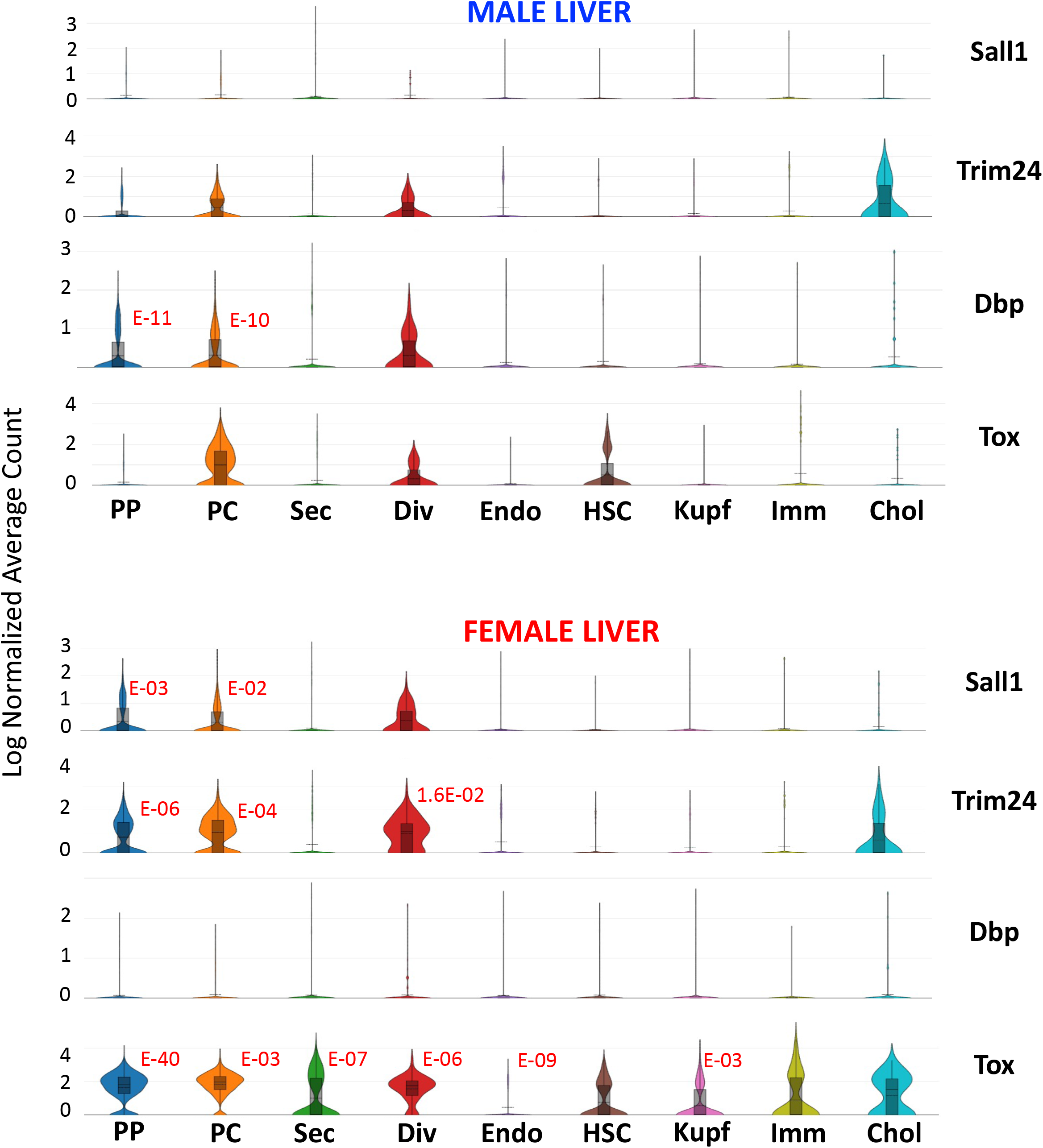
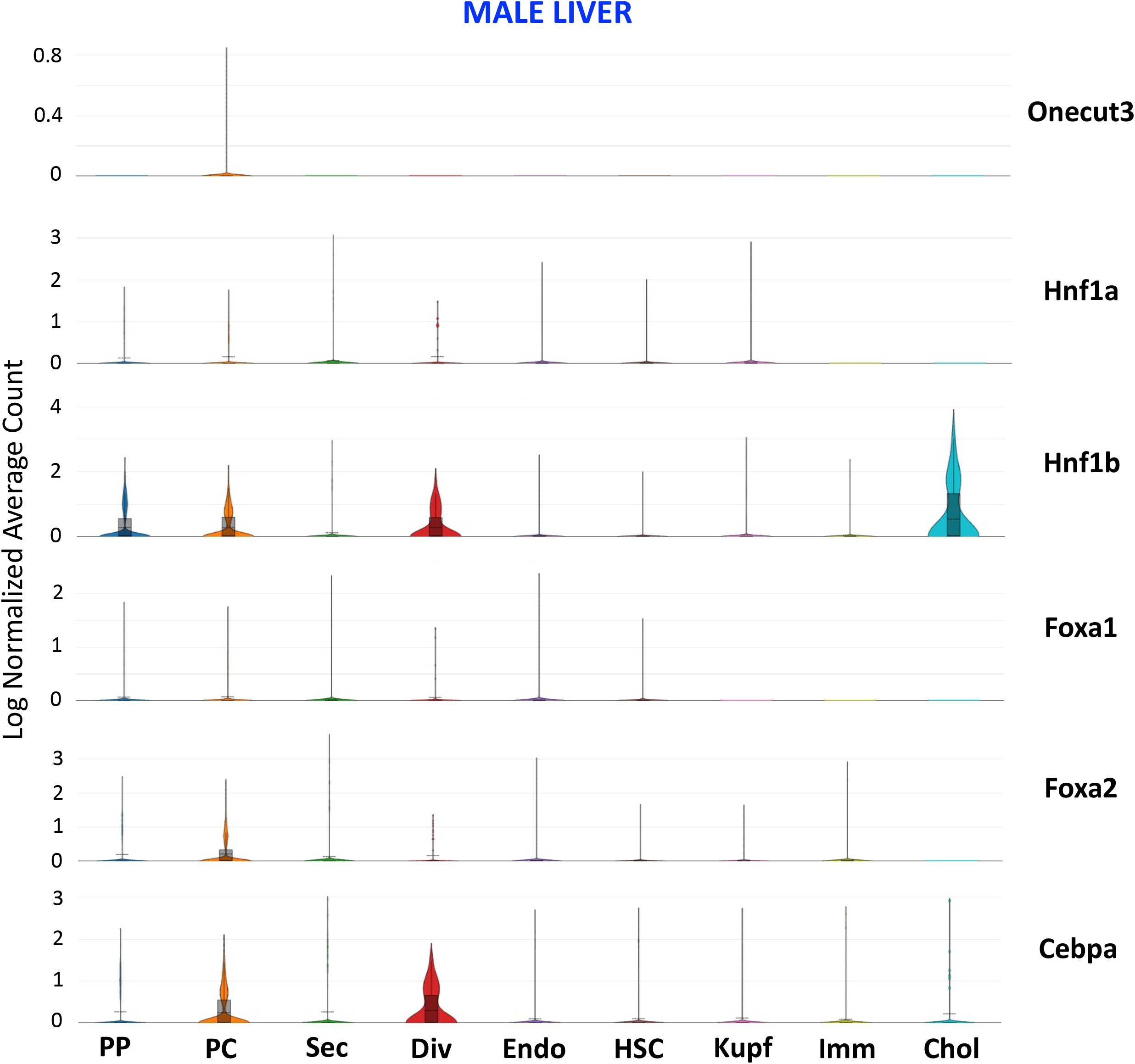
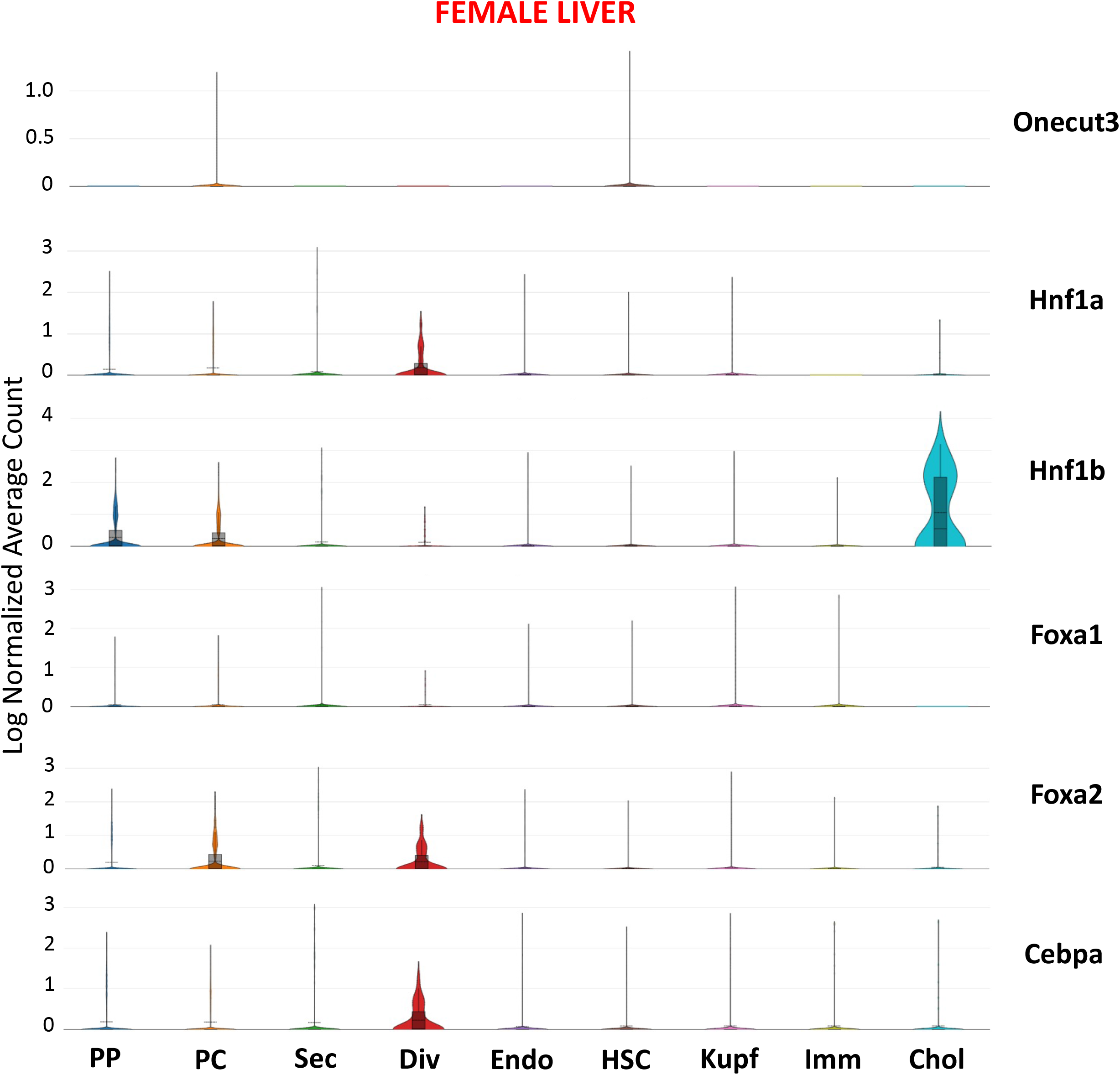
Expression of sex-biased genes and liver transcription factors. (**A, B**) Expression of key growth hormone signaling factors (Ghr, Jak2, Stat5b) and the feedback inhibitor of Grh/Jak2 signaling, Socs2, in hepatocytes vs NPCs, for both male (**A**) and female (**B**) liver. Mean expression values for Ghr are significantly higher in periportal (PP) and pericentral (PC) hepatocytes than in the five NPC clusters when combined for the analysis: Endothelial cells (Endo), Hepatic Stellate Cells (HSC), Kupffer cells (Kupf), Immune cells (Imm) and Cholangiocytes (Chol). Expression in also high in the dividing hepatocytes cluster (Div), but in the case of Stat5b is lower in secretory hepatocytes (Sec). Log normalized average expression based on UMI counts per nucleus for all nuclei in each cluster is indicated in black above the violin plot for each gene (see Methods). Significant differences in expression between [pericentral + periportal hepatocytes] versus [all 5 NPCs clusters] is indicated by the FDR values (red), based on analyses in **Table S2E**. NS, not significant. For Socs2 in female liver (**B**), values in purple, to the right of each violin plot, indicate the fold-change and FDR for significantly higher expression in female than male cells. (**C, D**) Violin plots indicating expression of liver transcription factors in each cluster from male and from female (control) livers. Many of these genes show elevated expression in periportal and pericentral hepatocytes and in dividing cells in male liver or in female liver compared to the 5 NPC clusters, with cholangiocytes being a notable exception for several of the transcription factors as indicated by the significance (FDR) values shown in red. Also note the sex differences in expression of Cux2, Bcl6, Sall1, Trim24, Socs2, and Tox, with significant FDR values indicated in red to the right of each violin plot (also see **Fig. 3**). Significance values are from **Table S2B**. (**F, G**) Transcription factors that show low expression in male (**F**) and female (**G**) liver.

**Fig. S3.**
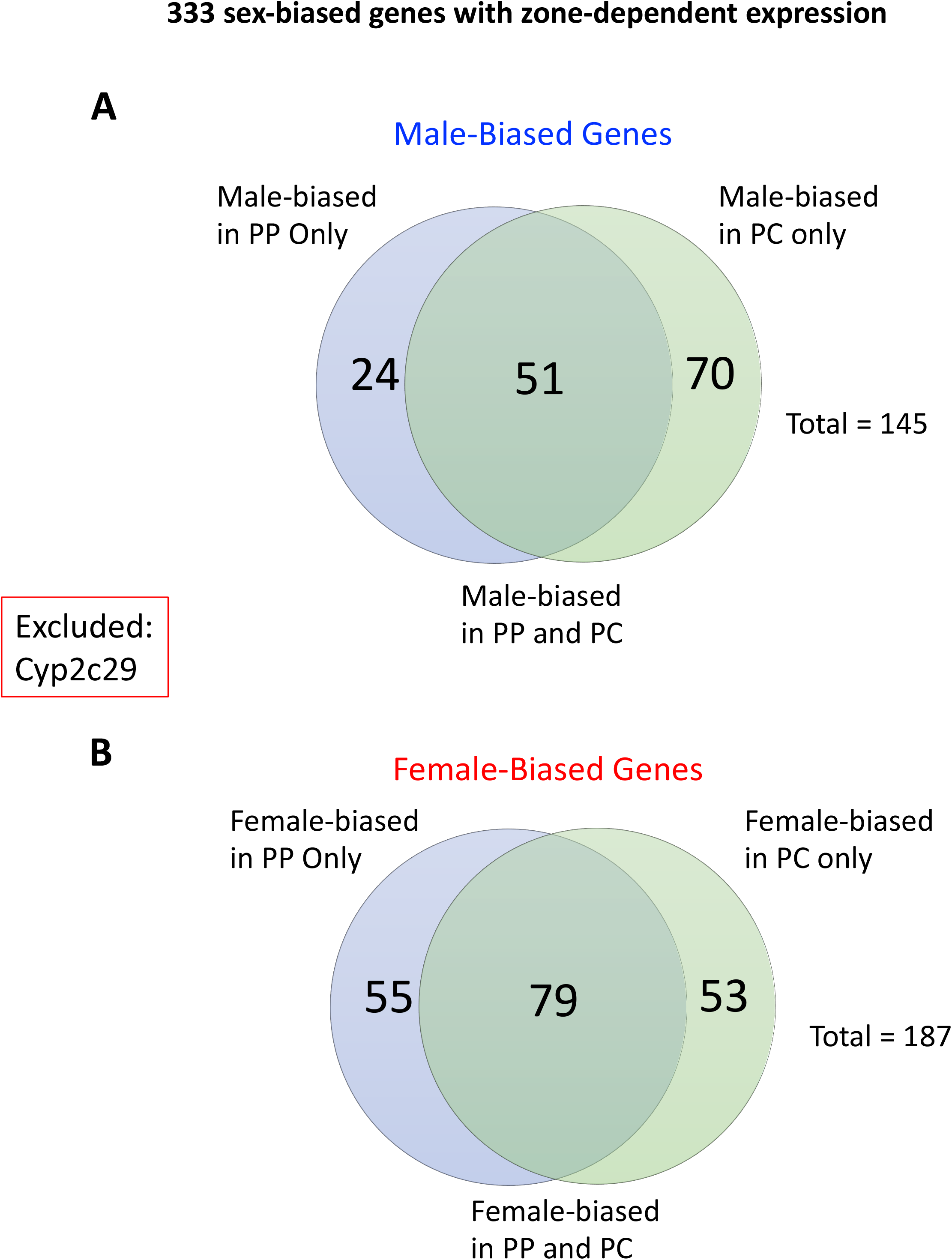
Set of 333 sex-biased genes that show sex-biased expression in only one hepatocyte zone. The number of male-biased (**A**) and female-biased (**B**) genes that are sex-biased in periportal (PP) hepatocytes, pericentral (PC) hepatocytes, or in both hepatocyte populations (overlap region). Full data is found in **Tables S3A and S3B**. Excluded is Cyp2c29, which shows a different sex-bias in PP hepatocytes (female-bias) than in PC hepatocytes (male-bias).

**Fig. S4.**
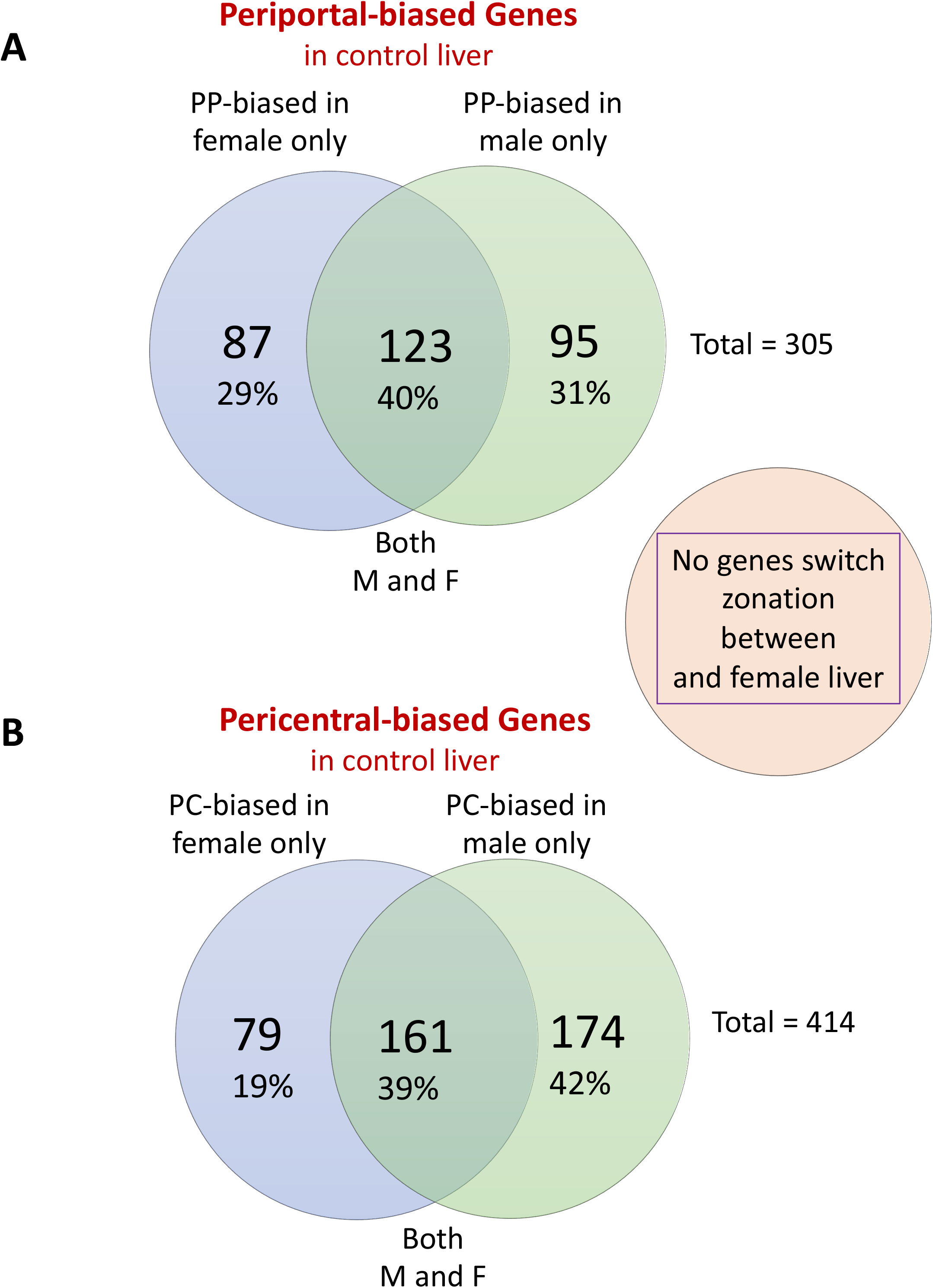
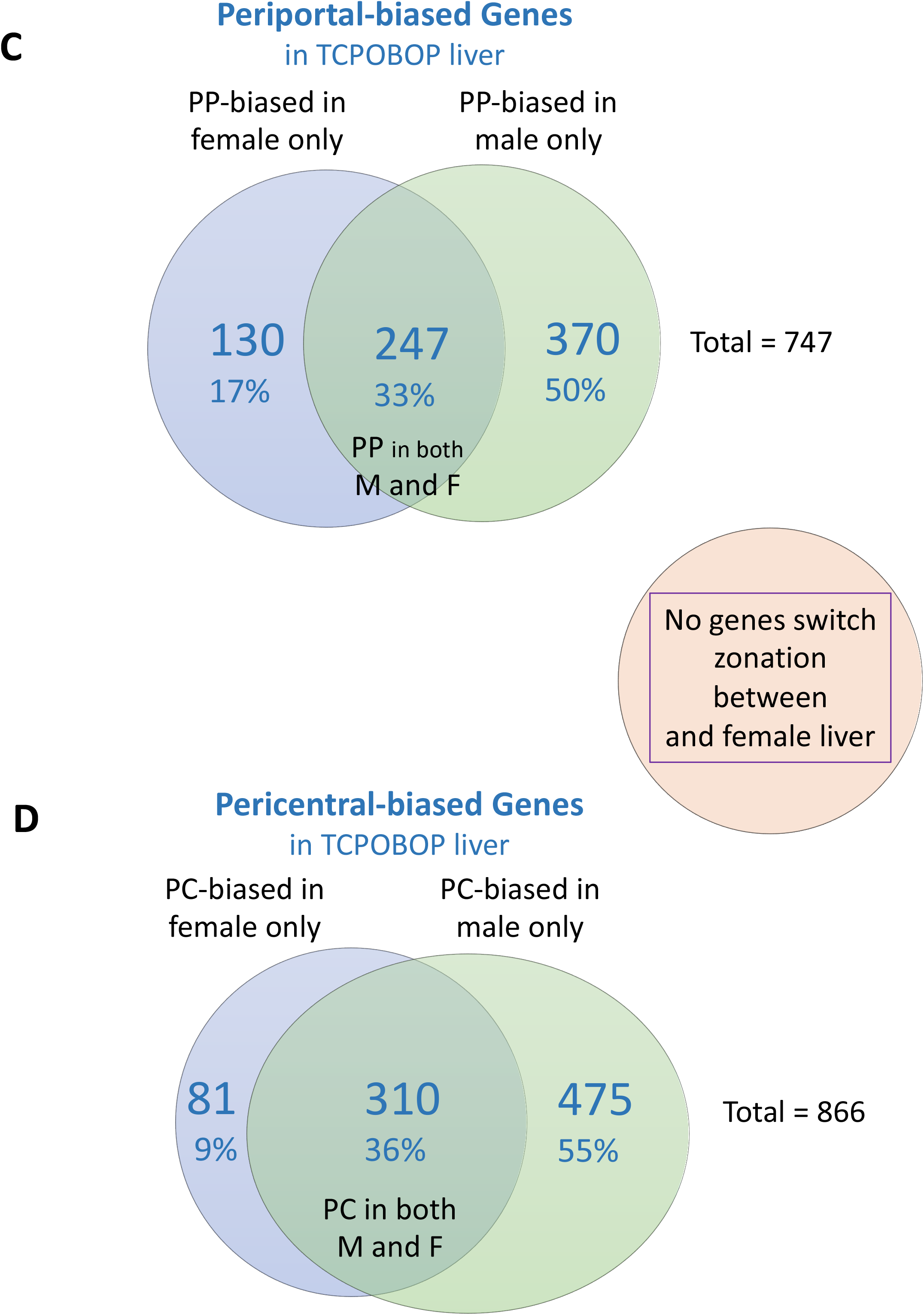
Number of zonation-biased genes that show zonated expression in control male or female liver (A, B), and in TCPOBOP-exposed male and female liver (C, D). Shown is the number of periportal-biased genes (**A, C**) and the number of pericentral-biased genes (**B, D**) that are significantly zonation-biased in male liver, female liver, or in both male and female liver (overlap regions). Full zonation-biased data for control and TCPOBOP-exposed hepatocytes is provided in **Tables S3A and S3B**. These data indicate that sex-dependent zonation patterns were more prevalent in TCPOBOP-exposed liver than in control liver (c.f., 60-61% of total zonated genes unique to either males or females in control liver vs 64-67% in TCPOBOP-exposed liver), consistent with the widespread effects of TCPOBOP on sex-biased gene expression reported in this study. Many more pericentral zonated genes were specific to males than were specific to females, in particular following TCPOBOP exposure (55% of all zonated genes unique to males versus 9% of all zonated genes unique to males in TCPOBOP liver, as compared to 42% in males versus 19% in females for control liver; panels D versus B).

**Fig. S5.**
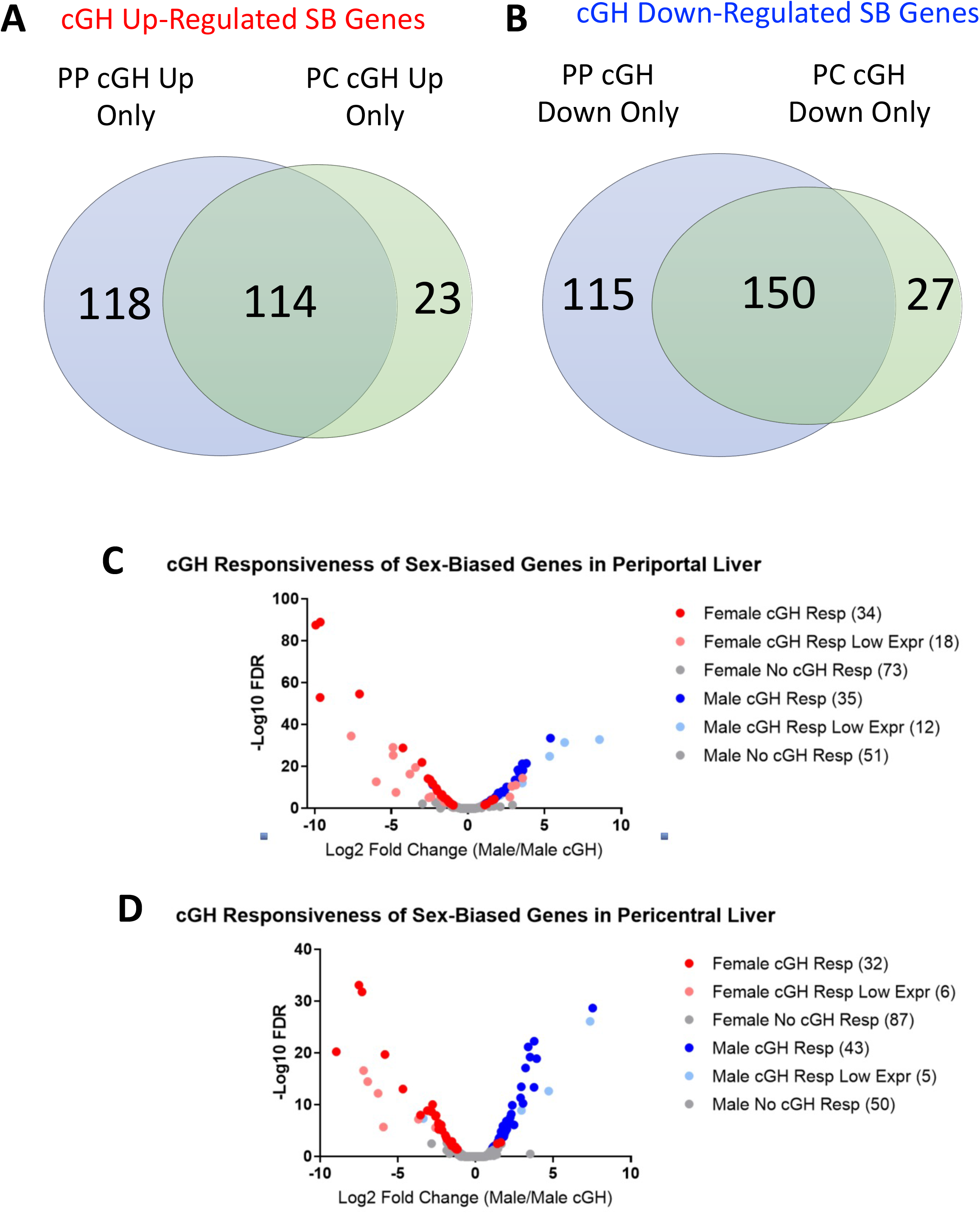
Sex-biased genes that respond to cGH in only one zone of the liver. Numbers of sex-biased (SB) genes that are up regulated (**A**) or down regulated (**B**) by cGH treatment in periportal (PP) hepatocytes, in pericentral (PC) hepatocytes, or in both hepatocyte populations (overlap region), based on Table S4B. (**C**, **D)**, Volcano plots showing the cGH responsiveness of 223 sex-biased genes whose maximum average expression intensity per cluster is >1 UMI in either periportal, pericentral or dividing hepatocyte clusters. Red, female-biased genes, which are largely up regulated in male liver following cGH treatment; blue, male-biased genes, which are primarily down regulated. Genes are colored as robustly responsive (dark red and dark blue), responsive but lowly expressed (light red and light blue), or not responsive (gray). Numbers in parenthesis: number of sex-biased genes in each group.

**Fig. S6.**
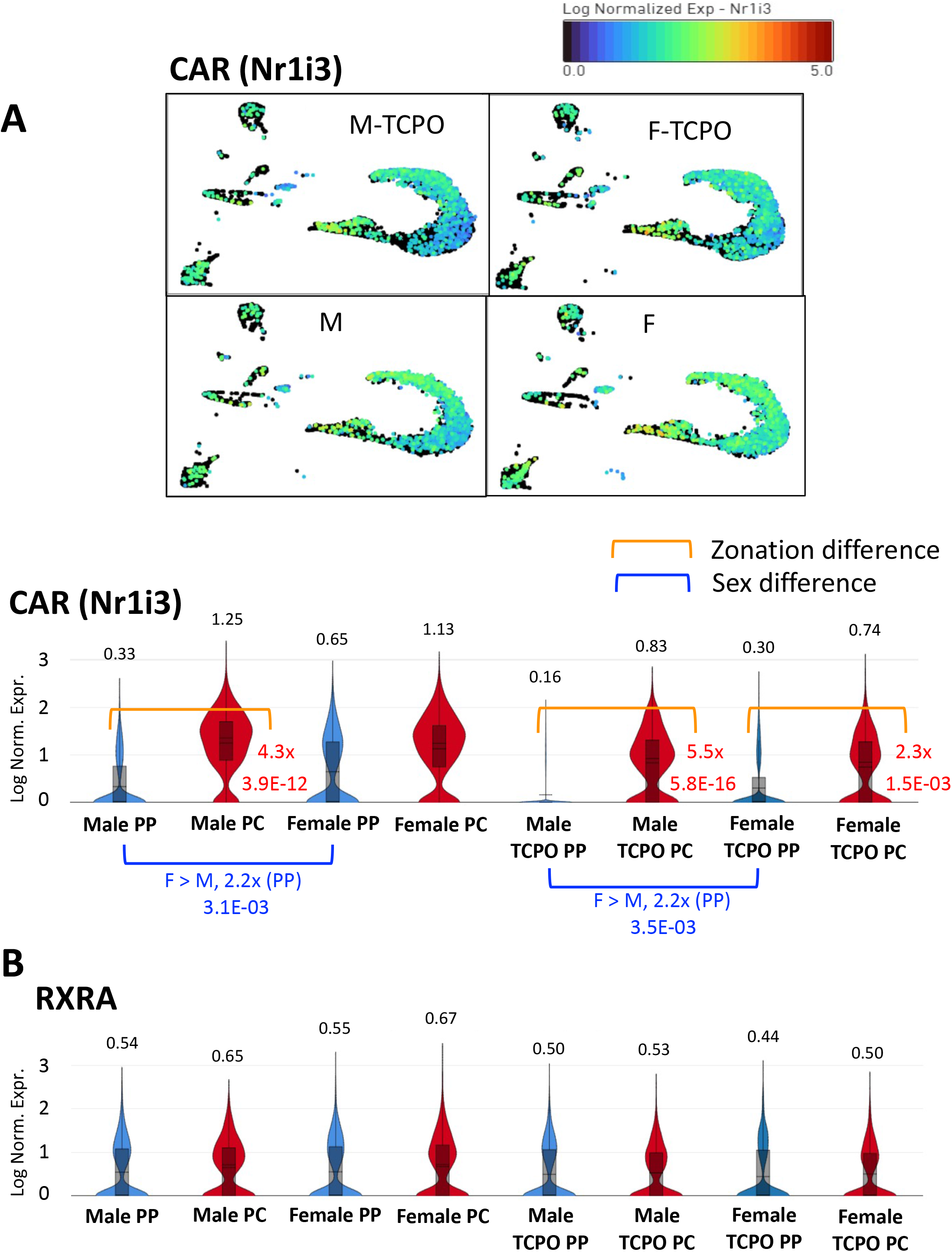
Zonation bias and sex differences for expression of Nuclear Receptors CAR and RXRA in control and in TCPOBOP-exposed hepatocytes. (**A**) CAR (gene symbol Nr1i3) shows significant pericentral-biased expression in male and female liver, with or without TCPOBOP treatment, as indicated. The four UMAPs (top) for CAR (gene symbol Nr1i3) present log normalized expression values for CAR in each nucleus for the four different groups. Violin plots in (**A**) and in (**B**) show log normalized average expression based on UMI counts per nucleus for all nuclei in the periportal and pericentral clusters for CAR and RXRA, respectively. The average expression value in each cluster is shown above the violin plots for each gene. Fold-change and FDR values (red) for zonation differences are shown to the right of the violin plot for those genes in each sex and treatment that show significant hepatocyte zonation-bias in their expression (from **Table S3A**), while sex differences are shown in blue below the violin plots (from **Table S5**). Consistent with the female predominance of CAR expression in periportal hepatocytes seen here, cGH infusion induced CAR expression in male hepatocytes, most notably in periportal hepatocytes (**Table S4A**).

**Fig. S7.**
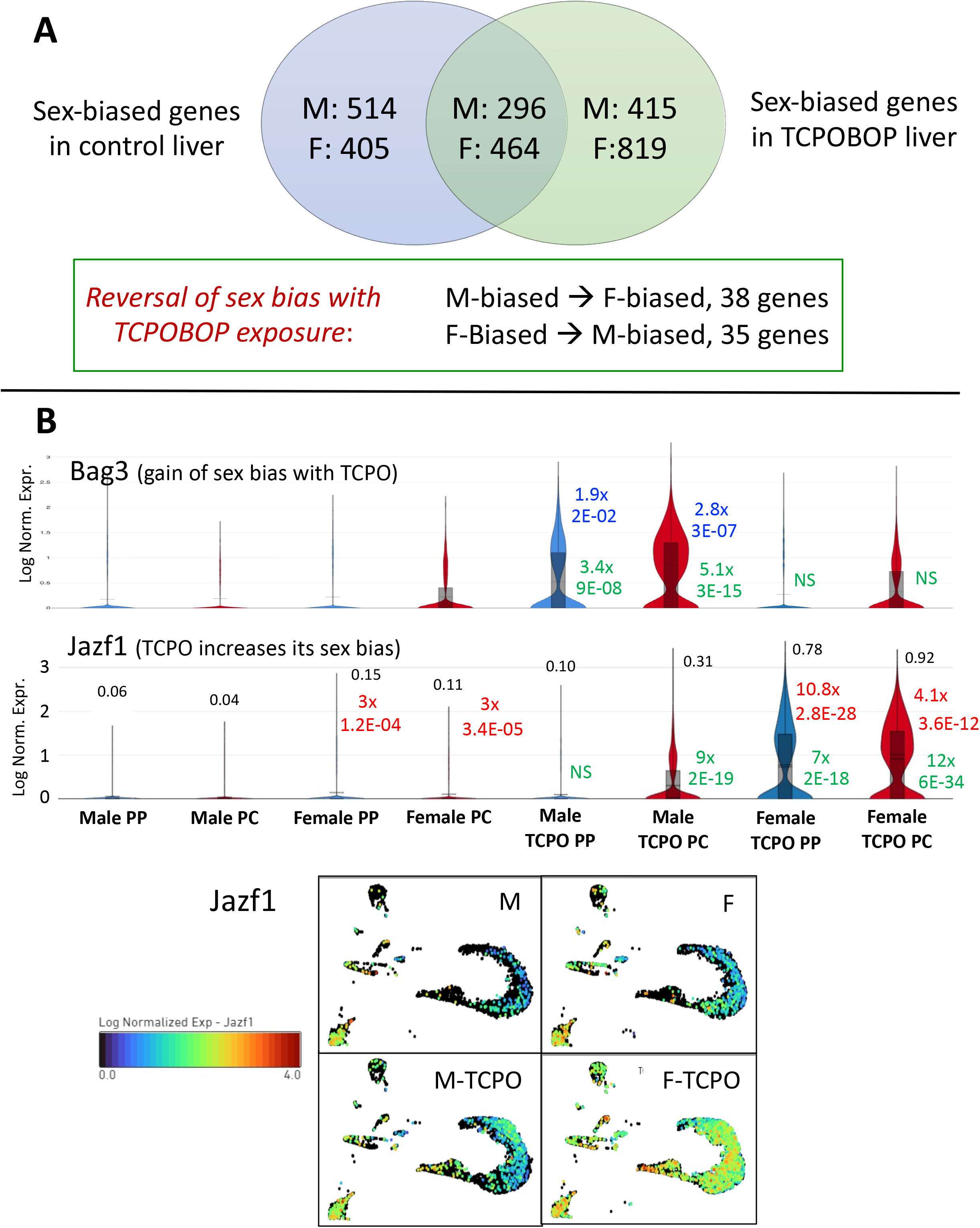
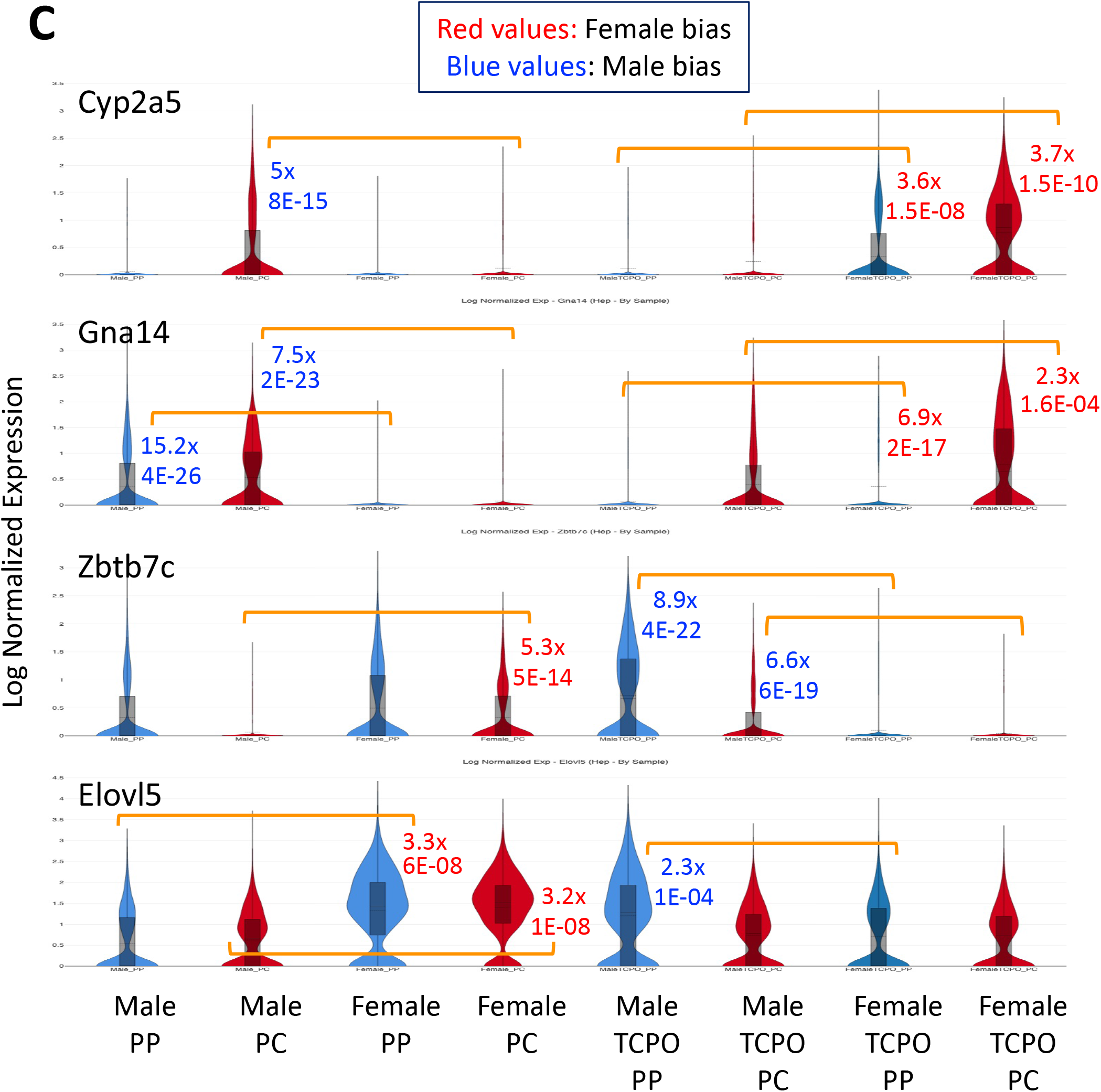
Impact of TCPOBOP on sex-biased genes in hepatocytes. (**A**) Venn diagram presenting the number of sex-biased genes identified in TCPOBOP-treated liver and their overlap with sex-biased genes identified in vehicle-treated liver (**Table S5**). A small number of sex-biased genes (38 and 35, as marked; 2.4% of all 2986 sex-biased genes) show the opposite site bias in control vs. TCPOBOP liver. (**B**) **Bag3** is a gene that gains male bias following TCPOBOP exposure, while **Jaz1** shows an increase in female-biased expression in TCPOBOP-exposed hepatocytes as compared to control hepatocytes. The four UMAP panels for Jazf1 at the bottom present log normalized expression in each nucleus in vehicle-treated and TCPOBOP-treated (‘TCPO’) male and female liver, as indicated. Violin plots present expression data as in Fig. 2. Fold change and FDR values to the right of the violin plots indicate significance of the male bias (blue values) or female bias (red values) in TCPOBOP hepatocytes, while the green values indicate responses to TCPOBOP exposure.

**Fig. S8.**
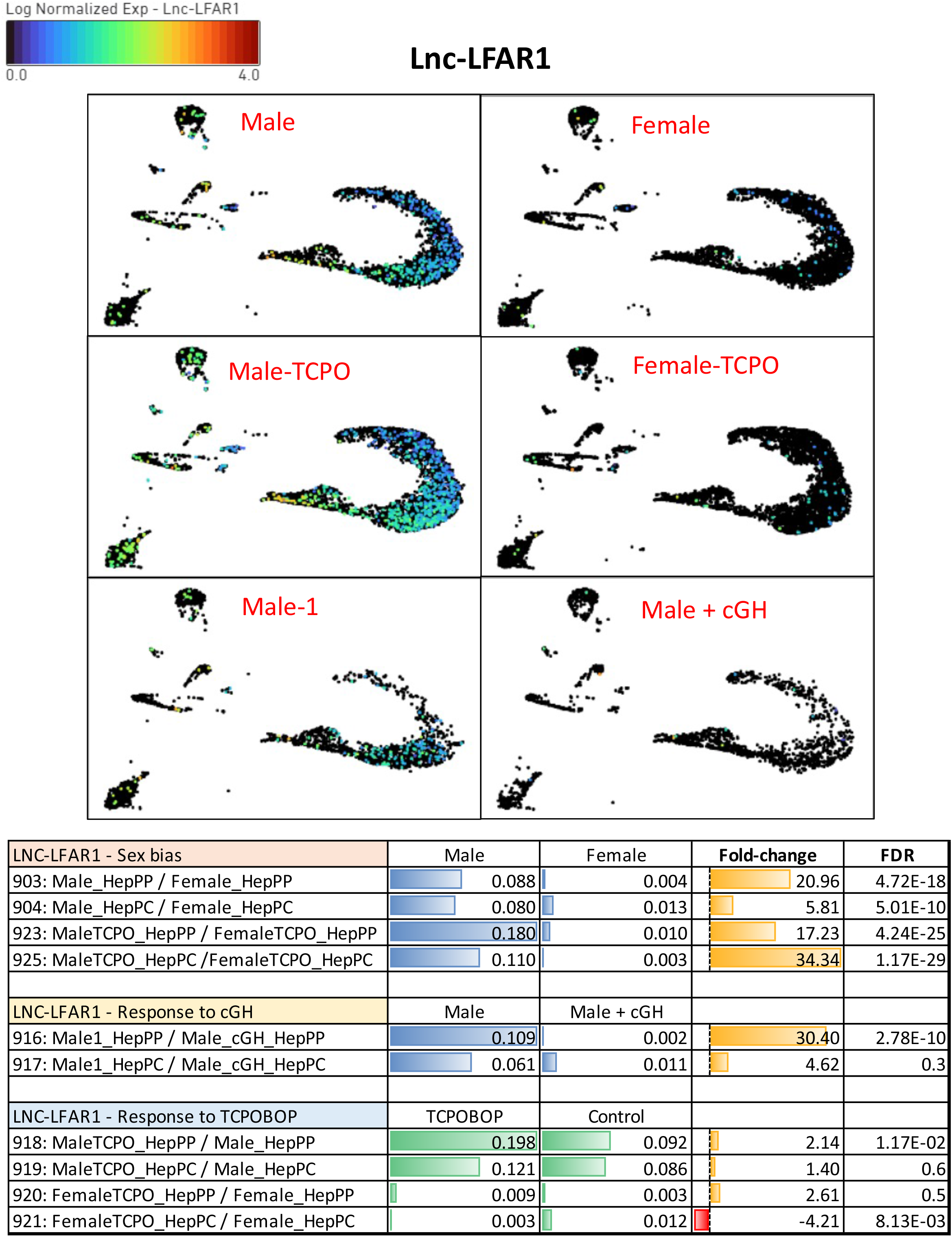
Expression of Lnc-LFAR1 across all six groups of liver samples. The six-panel UMAP indicates the log normalized expression of lnc-LFAR1 in each nucleus of each group, as per the color legend. This visualization reveals a pattern of male-specific expression, in both control and in TCPOBOP-exposed liver, and repression by cGH infusion (also consistent with the male-biased expression). Table below gives expression values and significance for control male and female liver, from Table S2B.

